# NRF2 pathway activation and SPP1⁺TREM2⁺ macrophages drive chemoradiotherapy resistance in esophageal squamous cell carcinoma

**DOI:** 10.64898/2026.02.20.706928

**Authors:** Jiaying Deng, Zhenzhen Xun, Hu Chen, Yanhong Cen, Yikai Luo, Wei Liu, Weiwei Chen, Junqiang Chen, Jinjun Ye, Xiansu Gao, Qingsong Fan, Jibin Song, Qi Chen, Yuan Li, Fei Ding, Menghong Sun, Yue Zhang, Ke Chen, Zhaozhen Zhang, Yiping He, Wenming Zhang, Jianqiang Liu, Zhuocheng Xin, Hongcheng Zhu, Qi Liu, Dashan Ai, Yun Chen, Jun Li, Kuaile Zhao, Han Liang

## Abstract

Esophageal squamous cell carcinoma (ESCC) is among the most aggressive cancers, with low rates of durable response to chemoradiotherapy and limited therapeutic options for relapsed disease. To uncover mechanisms of treatment resistance and relapse, we performed comprehensive multi-omics profiling of >100 pre-treatment and post-relapse ESCC tumors from a prospective clinical trial (NCT04694391), integrating whole-exome/genome sequencing, bulk RNA-sequencing, single-cell RNA sequencing, and spatial transcriptomics. We identify somatic alterations in *NFE2L2/KEAP1* in nearly 40% of relapsed patients, which are associated with upregulation of NRF2 signaling targets in resistant tumors and cell line models. At single-cell and spatial resolution, relapsed tumors are enriched for NRF2-activated epithelial cells that physically co-localize with immunosuppressive *SPP1⁺TREM2⁺* macrophages. This co-localization suggests a synergistic interaction between NRF2-driven tumor programs and macrophage-mediated immune suppression that promotes relapse after chemoradiotherapy. Our findings nominate *NFE2L2/KEAP1* mutations as predictive biomarkers for patient stratification and highlight therapeutic targeting of NRF2 signaling and *SPP1⁺TREM2⁺* macrophages as rational strategies to overcome resistance in ESCC.

## Introduction

Esophageal cancer ranks 11th in terms of incidence (510,716 new cases) and 7th in mortality overall (445,129 deaths) in 2022^1^. Of all cases, more than 50% occurred in China alone. Esophageal cancer can be categorized by two main histological subtypes: esophageal squamous cell carcinoma (ESCC) and adenocarcinoma (EAC). Most cases are ESCCs (84.8%) in global, 91.1% of esophageal cancer is ESCC in China^1^. Chemoradiotherapy is one of the most widely used approaches throughout the entire process of ESCC treatment, i.e., neoadjuvant chemoradiotherapy for operable ESCC^2^, definitive chemoradiotherapy for non-surgery locally advanced ESCC^3^, salvage chemoradiotherapy for recurrent local recurrence after definitive treatment^4^ and oligo-metastasis ESCC^5^, and palliative radiotherapy for advanced ESCC^6^. The risk of recurrence after neoadjuvant chemoradiotherapy and surgery remains high, especially among the 51 to 64% of patients who do not have a pathological complete response^2,4^. The incidence of local/regional failure and local/regional persistence of disease is 47% for locally advanced esophageal cancer treated with definitive chemoradiotherapy^3^. Therefore, investigating the molecular mechanisms underlying chemoradiotherapy resistance is urgently needed to develop more effective therapeutic strategies and improve clinical outcomes.

ESCC relapse after chemoradiotherapy can be attributed to both tumor-intrinsic factors and effects from the tumor microenvironment (TME)^7,8^. The tumor intrinsic factors include genetic intratumoral heterogeneity and clonal evolution, which drive adaptive resistance mechanisms^9^. On the other hand, TME-related factors such as hypoxia, immunomodulation, revascularization, and cancer-associated fibroblast-mediated ECM remodeling and fibrosis play a crucial role^10^. These changes in the TME are interconnected, with overlapping and widespread effects driven by key growth factors (TGFβ, VEGF, and PDGF), cytokines (TNF, CXCL12, IL1, IL2, IL6, and IL10), and transcription factors (NFκB and HIF1α)^11^. Despite this knowledge, it remains unclear how tumor-intrinsic heterogeneity interacts with the TME to drive ESCC relapse after chemoradiotherapy.

Applying cutting-edge multi-omics approaches to address this knowledge gap holds significant potential for unraveling the molecular basis of ESCC relapse and identifying novel therapeutic targets. Previous studies have predominantly relied on single-omics approaches, focusing on differences between primary tumors and adjacent normal tissues. These efforts have often been limited to creating basic ESCC tumor atlases^12–14^ or annotating the spatial distribution of specific stromal cell clusters^15^. While valuable, these observations underscore the complexity of the tumor TME) in primary ESCC and highlight the need for more comprehensive characterization in real-world settings with clear clinical response.

Here, we present a large-scale, multi-omics dataset generated from a clinical trial to investigate the factors driving chemoradiotherapy resistance in ESCC patients. Our analysis identifies mutations in *NFE2L2* and *KEAP1* as key drivers of resistance, by activating antioxidant response and promoting the recruitment of immunosuppressive *SPP1⁺TREM2⁺* macrophages. These findings highlight a potential predictive marker for identifying patients at risk of resistance to chemoradiotherapy. Furthermore, this study offers insights for the development of combination therapeutic strategies targeting the NRF2 pathway and *SPP1⁺TREM2⁺* macrophages with specific inhibitors, paving the way for more effective personalized treatments.

## Results

### Study Overview and Patient Characteristics

To rigorously investigate the molecular mechanisms underlying chemoradiotherapy resistance in ESCC patients, we conducted a multi-center, prospective clinical trial (NCT04694391) (**Figure 1A**) and recruited treatment-naïve patients with newly diagnosed ESCC. After undergoing chemoradiotherapy (or radiotherapy alone) and completing a minimum 2-year follow-up, patients were categorized into two groups: *Relapsed (Rel)*, defined as those experiencing local or metastatic relapse within two years, and *Non-Relapsed (NR)*, defined as those without such occurrences within three years (see **Methods**). Enrolled patients were further divided into three cohorts, each designed for different molecular profiling purposes: the discovery cohort, the mechanism investigation cohort, and the validation cohort.

**Figure 1.**
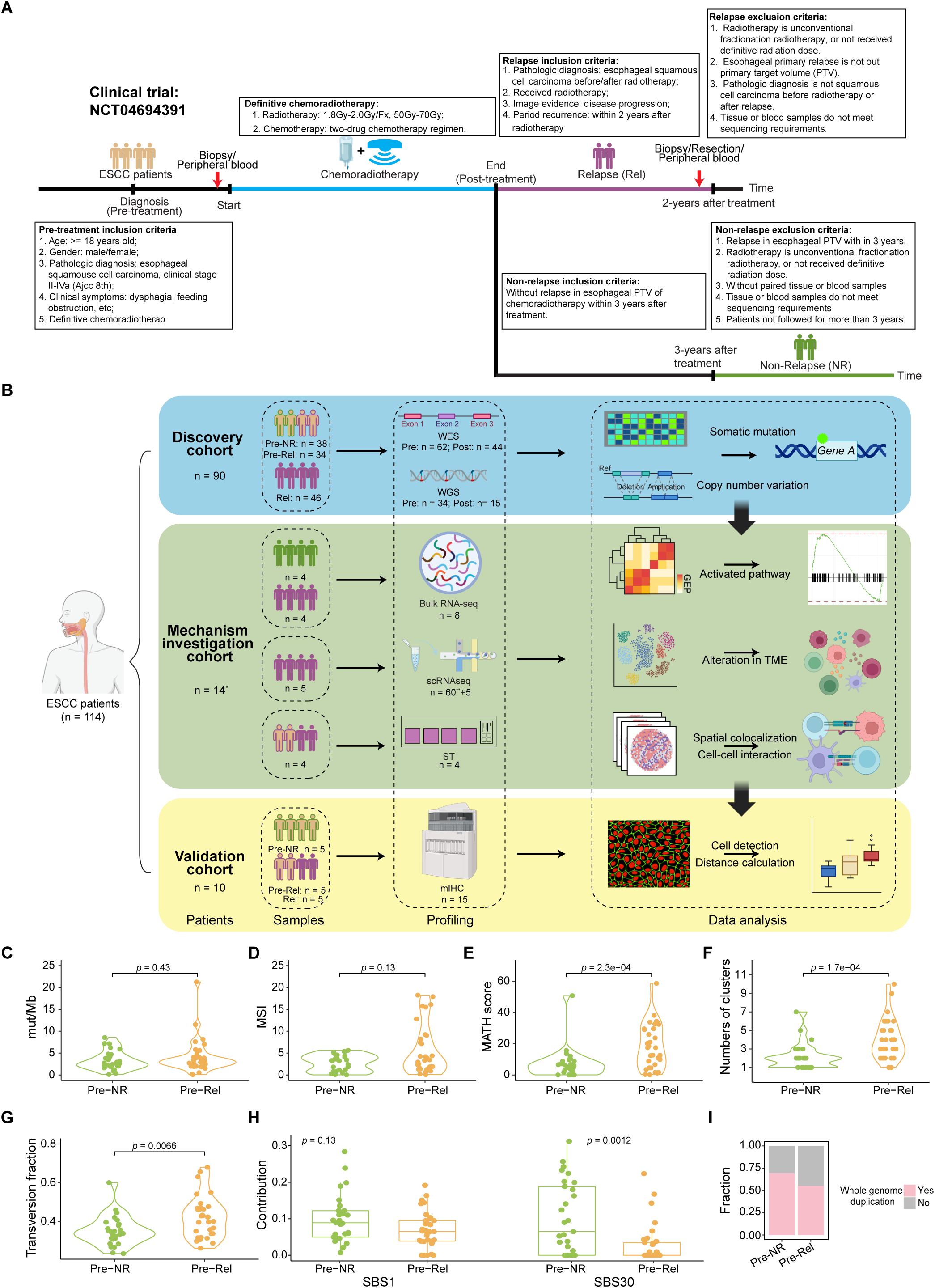
Study design and dataset overview. **A)** Flowchart illustrating patient enrollment and treatment timelines. **B)** Overview of sampling strategy for tumor biopsies, resection tissues, and blood samples across ESCC patients. Samples were processed for whole-exome/whole-genome sequencing (WES/WGS), bulk RNA-seq, single-cell RNA-seq, and spatial transcriptomics (ST) profiling. Numbers denote patient count and corresponding sample count. *One patient in the mechanism investigation cohort was sampled simultaneously for bulk RNA-seq and ST. **Includes 60 patients from a publicly available primary ESCC dataset. **C-G**) Violin plots displaying different features of tumor heterogeneity in pre-treatment samples, comparing relapsed and non-relapsed cases. This **H)** Box plots showing the contribution of different mutational signatures in pre-treatment samples, comparing relapsed and non-relapsed cases. The middle line in the box is the mean, the bottom and top of the box are the first and third quartiles, and the whiskers extend to the 1.5× interquartile range of the lower and the upper quartiles, respectively. **I)** Bar plot showing the fraction of whole-genome duplication (WGD) in pre-treatment samples, comparing relapsed and non-relapsed cases. (Discovery cohort with WGS: Pre-NR: n = 17; Pre-Rel: n = 17). C-H) P values are based on two-sided Wilcoxon rank sum tests. (Discovery cohort with WES: Pre-NR: n = 29; Pre-Rel: n = 33).

The discovery cohort aimed to systematically identify somatic alterations contributing to chemoradiotherapy resistance (**Figure 1B**, **Figure S1, Table S1**). Biopsy and peripheral blood samples were collected pre-treatment as the baseline, and post-treatment samples were obtained from relapsed patients whenever possible. This cohort included 90 patients: 38 in the NR group and 52 in the Rel group. Importantly, there were no significant differences in the distributions of age at diagnosis, gender, or tumor stages between NR and Rel patients in the cohort, ensuring an unbiased comparison in the subsequent analyses (**Figure S1C-E**). In terms of clinical samples obtained, 38 pre-treatment samples (Pre-NR) were collected from the NR group, and 34 pre-treatment samples (Pre-Rel) as well as 46 post-treatment relapsed samples (Rel) were collected from the Rel group. To identify key somatic point mutations, we profiled 106 tumor samples and their matched normal samples using whole-exome sequencing (WES), comprising 62 pre-treatment samples (29 Pre-NR and 33 Pre-Rel) and 44 Rel samples (**Figure S1A**). Among the Rel patients, 27 had paired pre- and post-treatment samples profiled with WES. To detect key somatic copy-number alterations (SCNAs), we profiled 49 tumor samples, including 34 pre-treatment samples (17 Pre-NR and 17 Pre-Rel) and 15 Rel samples, using whole-genome sequencing (WGS, **Figure S1B**).

The mechanism investigation cohort focused on assessing the effects of key somatic alterations on tumor-intrinsic pathways and the TME (**Figure 1B, Tables S2–S4**). This cohort included 14 patients analyzed using various molecular profiling strategies. Bulk RNA sequencing (RNA-seq) was performed on four Pre-NR and four Pre-Rel samples to identify altered signaling pathways associated with resistance (**Table S2**). Single-cell RNA sequencing (scRNA-seq) was used to profile five Rel samples, which were further integrated with a public dataset of 60 treatment-naïve ESCC samples to examine relapse-related effects on the TME (**Table S3**). To investigate relapse-related spatial interactions, matched pre- and post-relapse biopsies from two relapsed patients were profiled using 10× Visium spatial transcriptome sequencing (ST) (**Table S4**).

The validation cohort was designed to confirm findings from the discovery and mechanism investigation cohorts (**Figure 1B, Table S5**). Pre-treatment biopsy samples were collected from five NR patients, while both pre- and post-treatment biopsy samples were collected from five Rel patients. These samples were profiled using multiplex immunohistochemistry staining (mIHC) to validate the molecular and cellular features identified in the previous cohorts.

In total, this study included 114 patients and employed six profiling technologies, generating a comprehensive atlas of molecular features underlying chemoradiotherapy resistance in ESCC. By integrating tumor-intrinsic factors with alterations in the TME at both bulk and single-cell levels, this comprehensive resource enables us to elucidate how these factors collectively contribute to chemoradiotherapy resistance in ESCC.

### Pretreatment Relapsed Tumors Show Higher Intra-Tumoral Heterogeneity

To identify molecular features related to chemoradiotherapy resistance, we conducted a series of comparative analyses using Pre-NR and Pre-Rel samples from the discovery cohort. Pre-Rel samples did not show significant differences in tumor mutation burden (TMB) or microsatellite instability (MSI), two FDA-approved biomarkers, compared to Pre-NR samples (**Figure 1C, D**). We then assessed intra-tumoral heterogeneity using MATH^16^ (Mutant-Allele Tumor Heterogeneity) scores and the number of clonal/subclonal clusters (see **Methods**). The MATH score quantifies the distribution of variant allele frequencies of somatic mutations, with higher scores indicating greater intra-tumoral heterogeneity. Interestingly, Pre-Rel samples exhibited significantly higher intra-tumoral heterogeneity according to both metrics (**Figure 1E**, *p* = 2.3×10⁻⁴; **Figure 1F**, *p* = 1.7×10⁻⁴). These findings suggest that tumors with higher intra-tumoral heterogeneity are more likely to develop resistance, presumably due to an increased likelihood of harboring a persistent clone.

We next analyzed the composition of six possible base-pair substitutions and observed a significantly higher proportion of transversions in Pre-Rel samples (**Figure 1G**, *p* = 6.6×10⁻³). To further investigate the mutagenic processes driving these differential mutation contexts, we estimated the contributions of COSMIC mutation signatures (V3.2)^17^. Compared to Pre-NR samples, Pre-Rel samples exhibited a smaller contribution from Signature 30, which is associated with NTHL1-related base excision repair deficiency (**Figure 1H**, *p* = 1.2×10⁻³). In contrast, Signature 1, which is correlated with the age of cancer diagnosis and is one of the most prevalent and dominant signatures in cancer^18^, showed no significant difference, although there was a slightly higher contribution in Pre-NR samples (**Figure 1H**, *p* = 0.13). Finally, we examined differences in whole genome duplications (WGD) between Pre-NR and Pre-Rel samples. While Pre-Rel samples exhibited higher levels of aneuploidy (median ploidy 3.3 versus 2.4) and a greater fraction of patients with WGD, these differences did not reach statistical significance (**Figure 1I**). Together, these results suggest that while the mutational landscapes of Pre-NR and Pre-Rel tumors are broadly similar, Pre-Rel tumors display significantly higher intra-tumoral heterogeneity (**Figures 1E–F**). This highlights the need for further exploration of the molecular and cellular determinants driving chemoradiotherapy resistance.

### NFE2L2/KEAP1 Mutations are Key Determinants of Chemoradiotherapy Resistance

To identify specific genes contributing to chemoradiotherapy resistance, we compared the SCNA and mutation profiles of Pre-NR and Pre-Rel samples in the discovery cohort using WES and WGS data. Globally, no significant differences in deletions were observed between Pre-NR and Pre-Rel samples based on WGS-inferred SCNAs. However, Pre-Rel samples displayed significant amplification peaks in 2q31.2, 5p15.33, 17q21.31, 18p11.31, and 19p13.3 (**Figure 2A**). Overlaying previously reported ESCC driver genes onto these peaks revealed that *NFE2L2* and *HNRNPA3* were within the amplified 2q31.2 region. *NFE2L2* encodes the transcription factor NRF2, which regulates the cellular response to oxidative stress and has been implicated in resistance to chemo- or radiation therapy in multiple cancer types^19–25^. High-resolution SCNA analysis from WGS confirmed stronger amplification of *NFE2L2* in Pre-Rel samples, with 5 out of 17 Pre-Rel samples showing high-level amplification compared to none in the Pre-NR group (**Figure 2B**, *p* = 0.02).

**Figure 2.**
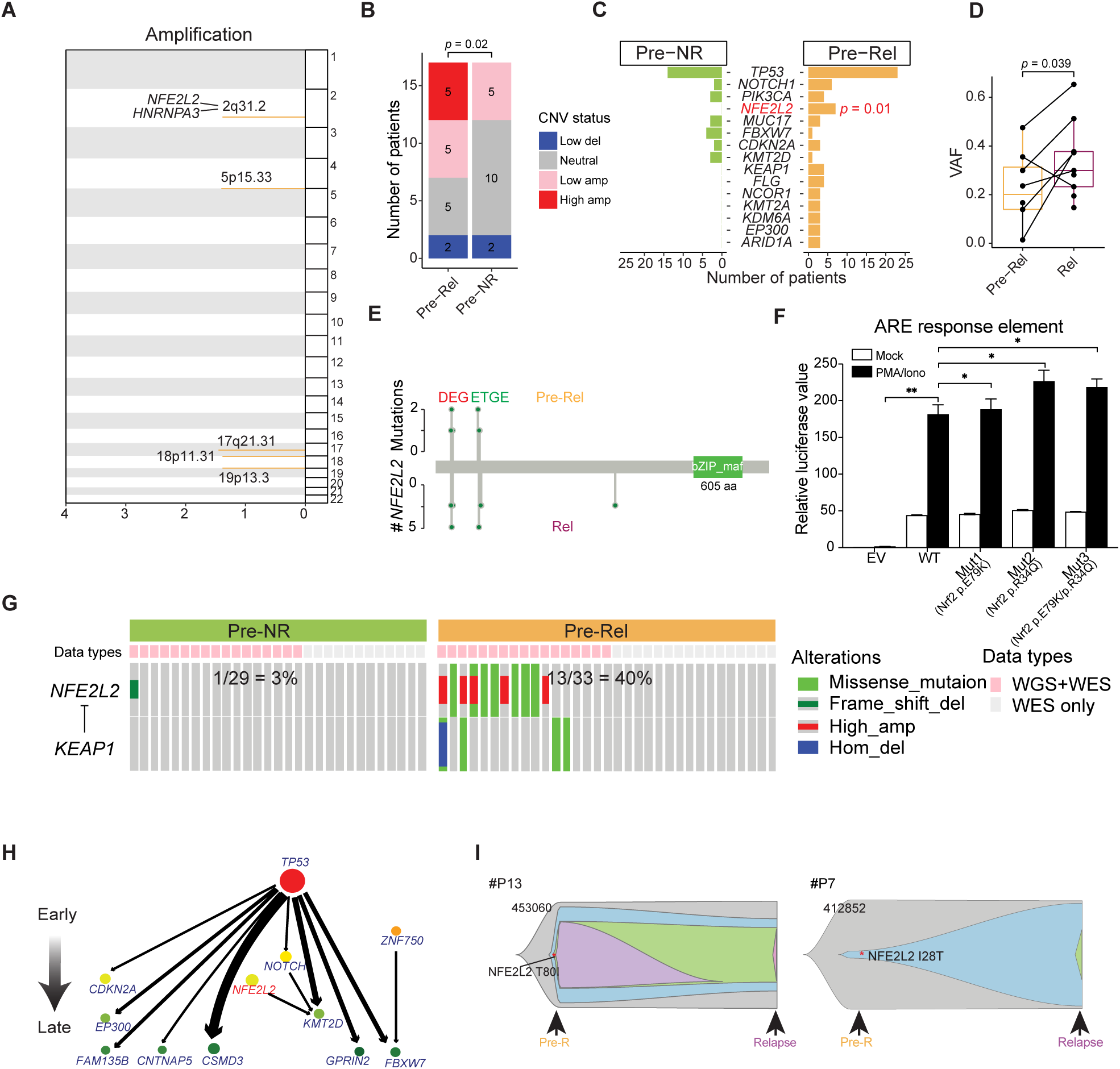
*NFE2L2* and *KEAP1* somatic alterations as major determinants of chemoradiotherapy resistance. **A)** Amplification plot identifying significant somatic copy number variation (SCNA) amplification peaks enriched in Pre-Rel samples. **B)** Bar plot displaying patient counts by SCNA status in pre-treatment samples, comparing relapsed and non-relapsed cases. P-value is based on chi-squared test. A-B) Discovery cohort with WGS data: Pre-NR: n = 17; Pre-Rel: n = 17. **C)** Bar plot showing the top-ranked mutations, highlighting their differential enrichment in relapsed pre-treatment samples. P-value is based on Fisher exact test. (Discovery cohort with WES: Pre-NR: n = 29; Pre-Rel: n = 33). **D)** Box plot comparing the variant allele frequencies (VAF) of *NFE2L2* mutations in matched pre-and post-chemoradiotherapy of relapsed patients. P-value is based on paired Wilcoxon rank test. Discovery cohort: 7 pairs with NFE2L2 mutations either in Pre-Rel or Rel samples. **E)** Mapping of *NFE2L2* mutations to two functional domains: DEG and ETGE, revealing their enrichment. Only missense mutations are shown. **F)** Bar plot illustrating differences in ARE (antioxidant response element) activity across varying *NFE2L2* mutation conditions. P-values are based on t-test. *, P < 0.05; **, P < 0.01. **G)** Comparative bar plots showing *NFE2L2* and *KEAP1* alterations in pre-treatment samples of non-relapsed (left) and relapsed (right) patients. Discovery cohort with WES: Pre-NR: n = 29; Pre-Rel: n = 33; WGS data are incorporated whenever possible. **H)** Tumor Evolutionary Directed Graph of paired samples depicting potential evolutionary trajectories of ESCC under chemoradiotherapy. This analysis is based on 27 paired Pre-Rel and Rel samples with WES data in the discovery cohort. **I)** Representative clonal evolution trajectories from two ESCC patients, highlighting clonal mutations of *NFE2L2*. Different clones are shown in different colors.

We next assessed the mutation frequency of significantly mutated genes (SMGs) reported in published ESCC studies^26,27^. The most frequently mutated genes in this study were *TP53*, *NOTCH1*, *PIK3CA, NFE2L2, MUC17, FBXW7, CDKN2A, KMT2D, KEAP1,* and *FLG* (**Figure 2C**). Among them, *NFE2L2* was the only gene showing a significant difference between Pre-Rel and Pre-NR samples, with mutations present in 7 Pre-Rel samples and absent in Pre-NR samples (**Figure 2C**, *p* = 0.01). We hypothesized that mutations conferring survival advantages against chemoradiation would be further enriched in Rel samples. Consistent with this, we observed significantly higher variant allele frequencies (VAFs) for *NFE2L2* mutations in Rel samples compared to their matched Pre-Rel samples (**Figure 2D**, *p* = 0.039). These mutations clustered within the DEG and ETGE functional domains of *NFE2L2*, where its negative regulator KEAP1 binds, suggesting that these mutations likely activate N*FE2L2* (**Figure 2E**). Functional assays further demonstrated that *NFE2L2* mutations led to significantly elevated antioxidant response element (ARE) activity under PMA/ionomycin treatment (**Figure 2F, Figure S2**).

Given that NFE2L2 and KEAP1 are central regulators of the NRF2 signaling pathway, we examined the combined mutational profiles of these genes in pre-treatment samples using WES, integrating point mutation, small indel, and SCNA data. Alterations in *NFE2L2* or *KEAP1* were significantly more frequent in Pre-Rel samples (13/33, 40%) than in Pre-NR samples (1/29, 3%) (**Figure 2G**, *p* = 7×10⁻⁴).

To gain insights into the role of *NFE2L2* mutations in ESCC tumor relapse, we analyzed potential evolutionary trajectories. Tumor evolutionary directed graphs (TEDG)^28^ constructed for 27 paired Pre-Rel and Rel samples revealed that *TP53* mutations were early events, while mutations in *NOTCH1, NFE2L2, CDKN2A, ZNF750, EP300*, and *KMT2D* accumulated later. Mutations in *FAM135B, CNTNAP5, CSMD3, GPRIN2*, and *FBXW7* represented the latest events, with *FBXW7* being a more recurrence-exclusive late event (**Figure 2H**). Most ESCC cases followed a branched evolutionary trajectory, consistent with the module splice model of tumor evolution. Clonal evolution trajectories from two exemplary patients highlighted how clonal mutations (e.g., *NFE2L2* T80I in patient #13) and subclonal mutations (e.g., *NFE2L2* I28T in patient #7) likely conferred resistance. The surviving clones or subclones subsequently acquired additional mutations, increasing tumor heterogeneity (**Figure 2I**).

Collectively, these findings identify *NFE2L2* and *KEAP1* somatic alterations, particularly gain-of-function mutations in *NFE2L2*, as major molecular drivers of chemoradiotherapy resistance in ESCC. Notably, aberrations in these two genes alone explain nearly half of early recurrences, underscoring their potential as predictive biomarkers for selecting patients most likely to benefit from chemoradiotherapy.

### NRF2 Pathway Activation Correlates with Resistance to Chemoradiotherapy

The above analysis suggests that mutations in these critical regulators of NRF2 signaling result in constitutive NRF2 activation, which in turn drives the transcription of ARE-mediated genes, thereby conferring resistance to therapy (**Figure 3A**). However, NRF2 activation may occur independently of mutations in *NFE2L2* and *KEAP1*. Notably, a substantial proportion of patient samples without these somatic alterations also exhibit NRF2 signaling activation, as observed in the TCGA ESCC cohort (**Figure S3**). To investigate the role of NRF2 signaling activation in chemoradiotherapy resistance, we focused on the mechanism investigation cohort. Leveraging bulk RNA-seq data from four Pre-NR and four Pre-Rel samples, we identified differentially expressed genes between these two groups. The analysis revealed a significant upregulation of the NRF2 pathway in Pre-Rel samples (**Figure 3B**, *p* < 0.001), with marked upregulation of downstream NRF2 target genes in Pre-Rel samples compared to Pre-NR samples (**Figure 3C**).

**Figure 3.**
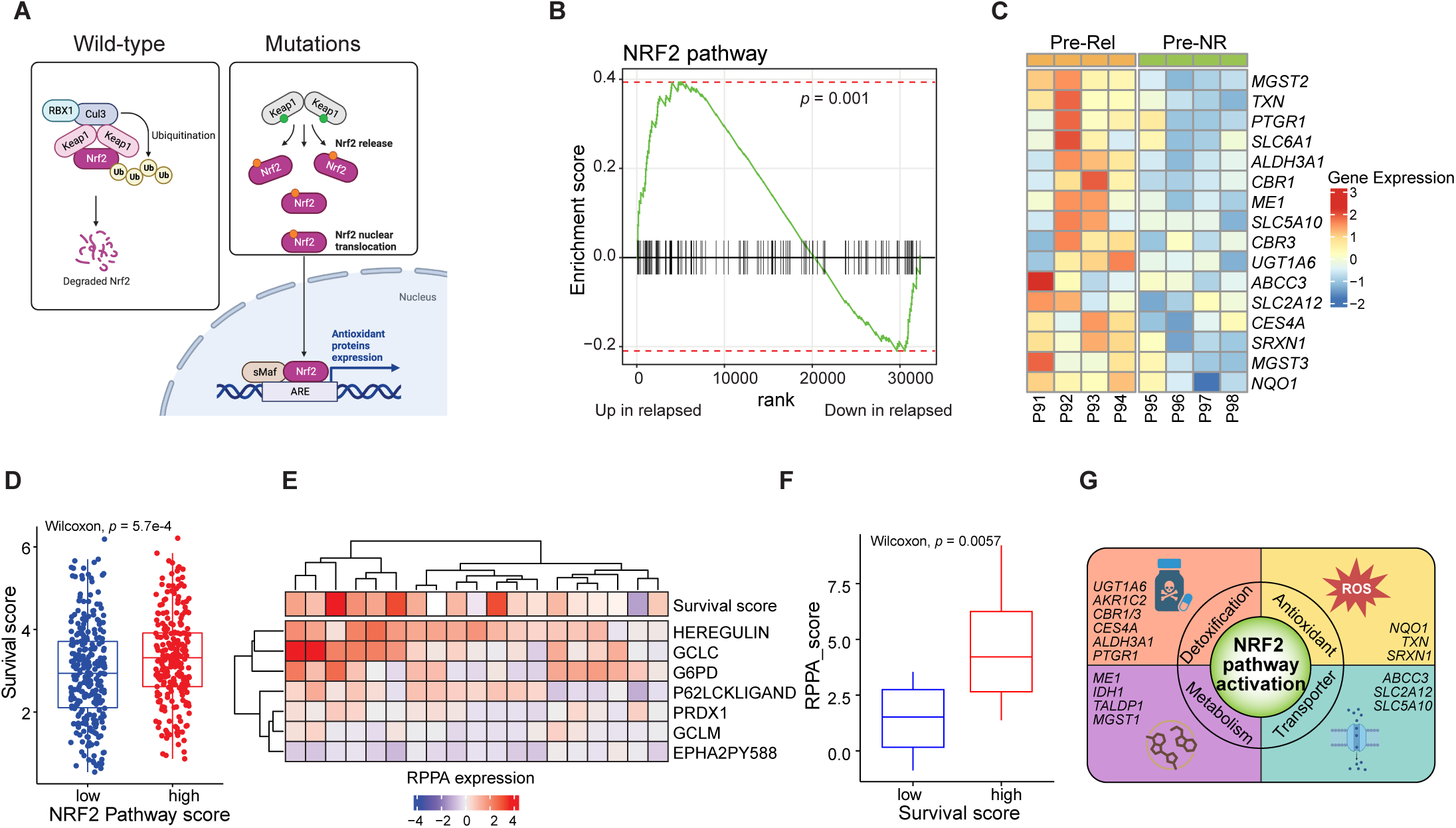
NRF2 pathway activation contributes to ESCC relapse. **A)** Schematic diagram illustrating how mutations in the NRF2 pathway (*NFE2L2* and *KEAP1*) result in enhanced antioxidant response by activating NRF2 target genes. **B)** Gene set enrichment analysis showing significant upregulation of the NRF2 pathway in pre-treatment samples of relapsed cases (Pre-Rel) compared to those of non-relapsed (Pre-NR) cases. The mechanism investigation cohort with bulk RNA-seq data: Pre-NR: n = 4; Pre-Rel: n = 4. **C)** Heatmap demonstrating the upregulation of key NRF2 pathway components in Pre-Rel samples relative to Pre-NR samples. **D)** Boxplot illustrating cancer cell survival rates, stratified by NRF2 pathway activation scores. Cell lines were classified equally into two groups based on the NRF2 pathway activation scores. P value is based on two-sided Wilcoxon rank sum tests. **E)** Heatmap of NRF2 target protein expression measured using reverse-phase protein arrays (RPPA) across 19 cancer cell lines. **F)** Comparison of the total expression of NRF2 target proteins between cell lines with low (AUC_low) and high (AUC_high) survival under radiation. P value is based on two-sided Wilcoxon rank sum tests. **G)** Functional categorization of NRF2 pathway downstream target genes upregulated in Pre-Rel samples. The central circle depicts NRF2 pathway activation. Surrounding it, an outer segmented ring highlights four functional categories commonly associated with NRF2 downstream targets: detoxification, antioxidant, metabolism, and transporter. Each colored quadrant represents one of these categories and displays the corresponding NRF2-regulated genes upregulated in the Pre-Rel samples, illustrating the functional diversity of NRF2-mediated transcriptional programs.

To validate the functional impact of NRF2 pathway activation independently, we utilized a panel of cancer cell lines with quantitatively measured survival under irradiation^29^. A significant positive correlation was observed between the NRF2 pathway gene-expression scores and survival scores across all cancer cell lines (**Figure 3D**, *p* = 5.7×10⁻⁴). We further confirmed this relationship at the protein level using reverse phase protein array (RPPAs) analysis of 19 ESCC cell lines^30^. The overall RPPA scores for NRF2 pathway downstream targets were significantly higher in cell lines with elevated survival scores compared to those with lower survival scores (**Figure 3F**, *p* = 5.7×10⁻³).

We next investigated the molecular mechanisms through which NRF2 pathway activation contributes to chemoradiotherapy resistance, focusing on a subset of well-characterized NRF2 downstream targets previously implicated in treatment resistance. Notably, we observed significant upregulation of these targets across four key cellular processes (**Figure 3G**) as reported in the literature^31,32^. These include detoxification enzymes (*UGT1A6*^33^, *CBR1/3*^33^, *CES4A, ALDH3A1*^34^, *and PTGR1*^33^), antioxidant defense genes (*NQO1*^35^, *TXN*^35^, *and SRXN1*^35^), metabolic regulators (*ME1*^36^, *IDH1*, *TALDP1*, and *MGST1*^33^), and transporters (*ABCC3*^37^, *SLC2A12*^34^, and *SLC5A10*^34^). These findings suggest that NRF2-mediated transcriptional reprogramming enhances cellular detoxification, redox balance, metabolic flexibility, and drug efflux capacity, collectively contributing to therapy resistance.

These findings from patient samples and cell lines highlight the pivotal role of NRF2 pathway activation in enhancing tumor-intrinsic resilience and survival under DNA-damaging treatment. This mechanistic insight underscores NRF2 signaling as a potential target for overcoming chemoradiotherapy resistance.

### Enrichment of NRF2-Activated Tumor Cells and *SPP1*^+^*TREM2*^+^ Macrophages in Relapsed Samples

To investigate the relationship between NRF2 signaling activation in tumor cells and its impact on the TME, we analyzed scRNA-seq data from five Rel samples in the mechanism investigation cohort. For comparison, we incorporated a publicly available ESCC scRNA-seq dataset^12^ of treatment-naive primary tumors and adjacent normal tissues from 60 patients (**Figure 1B**). Following rigorous quality control and data integration using Harmony^38^ (see **Methods**), our analysis encompassed 240,634 cells from relapsed and treatment-naïve tumors of 65 patients. Cell-type annotation based on marker gene expression identified major cell populations, including epithelial cells, myeloid cells, T and NK cells, B and plasma cells, endothelial cells, and fibroblasts (**Figure S4A-C**).

Further investigation of epithelial cell heterogeneity identified nine distinct subtypes using cNMF^39^ and based on subtype-specific markers (**Figure 4A, B**). Among these, a subset of epithelial cells exhibited high expression of *AKR1B10*, a marker of NRF2 pathway activation^40^. Using NRF2 pathway scores derived from MsigDB^41^ and a published NRF2 activation signature score^42^, this *AKR1B10*-high subtype displayed the highest NRF2 activation levels (therefore termed as *Epi_NRF-activated*) (**Figure 4C, Figure S4D**). Functional enrichment analysis revealed its associations with cellular detoxification, glutathione metabolism, and the *KEAP1-NFE2L2* pathway (**Figure S4E**). Strikingly, the proportion of NRF2-activated epithelial cells increased progressively from normal tissues to treatment-naive tumors and further to relapsed tumors (**Figure 4D**, **Figure S4F**). These findings were validated using mIHC on the independent validation cohort, confirming an enrichment of AKR1B10-positive and PanCK-positive cells in Rel samples (**Figure 4E**). The results underscore the role of NRF2-activated tumor cells in promoting chemoradiotherapy resistance.

**Figure 4.**
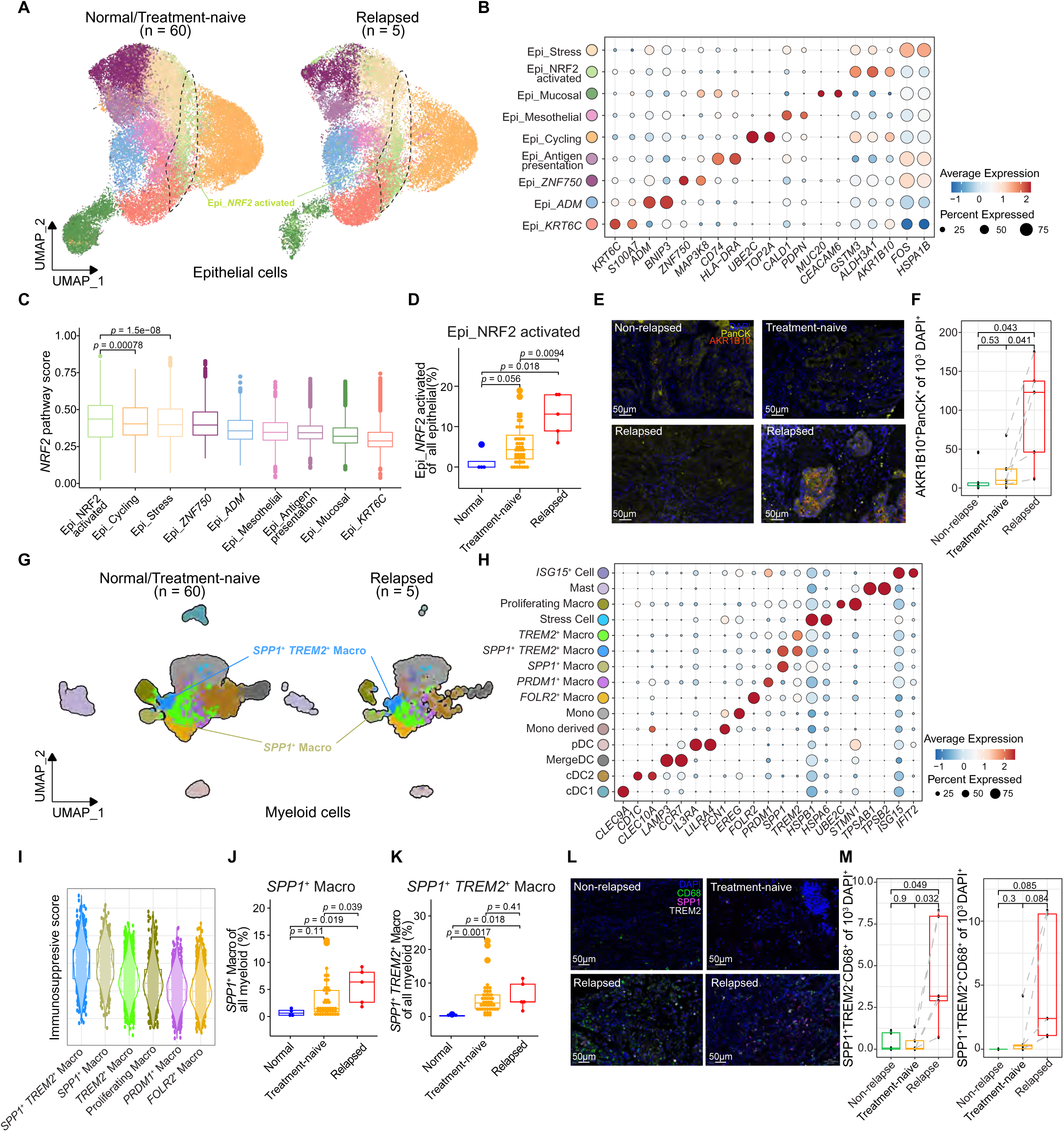
Enrichment of NRF2-activated epithelial cells and SPP1⁺TREM2⁺ macrophages in relapsed ESCC samples. **A, G**) UMAP embeddings of epithelial cells (A) and myeloid cells (G), color-coded by annotated cell states **B, H)** Dot plots showing normalized gene expression used for cell state annotations in epithelial cells (B) and myeloid cells (H). Dot size represents the proportion of expressing cells in each cluster, and color indicates expression intensity. **C)** Boxplot illustrating differences in NRF2 pathway activation scores across epithelial cell clusters. **D, J, K**) Bar plots showing the proportions of NRF2-activated epithelial cells (D) within all epithelial cells, SPP1⁺ macrophages (J), and SPP1⁺TREM2⁺ macrophages (K) within all myeloid cells across adjacent normal, primary, and relapsed tumor tissues. **E, L**) Representative multiplex immunohistochemistry (mIHC) images of pre-treatment non-relapsed (NR) tissues, and paired pre- and post-treatment relapsed tissues showing PanCK⁺AKR1B10⁺CD68⁻ cells (E) and SPP1⁺TREM2⁺CD68⁺ cells (L). Scale bars: 50 µm. **F, M**) Boxplots comparing the normalized counts of AKR1B10⁺PanCK⁺ cells (F), SPP1⁺TREM2⁻CD68⁺ cells (M, left), and SPP1⁺TREM2⁺CD68⁺ cells (M, right) across pre-treatment NR (green, n = 5), paired pre-treatment (orange, n = 5), and post-treatment (red, n = 5) relapsed tissues in the validation cohort. **I**) Boxplot illustrating differences in immunosuppressive scores across macrophage sub-clusters. UMAP, uniform manifold approximation and projection. For boxplots, the middle line represents the mean; the box edges indicate the first and third quartiles; whiskers extend to 1.5× the interquartile range. P values are calculated using a two-sided Wilcoxon signed-rank test in (C), (D), (F), (J), (K), and (M). (A-C) and (G-I) are based on the scRNA-seq data (n = 65) of the mechanism investigation cohort that combined the five Rel samples profiled in this study and a public dataset of 60 treatment-naïve ESCC samples. For fair comparison, stage I tumor samples in the public dataset were excluded.

We next examined myeloid cells, a major component of the TME, to understand their contribution to relapse in ESCC. We identified 15 myeloid cell subtypes, including six macrophage subtypes (*FOLR2^+^, PRDM1^+^, SPP1^+^, TREM2^+^, SPP1^+^TREM2^+^,* and proliferative macrophages), two monocyte subtypes (monocytes and mono-derived cells), four dendritic cell subtypes (cDC1, cDC2, mergeDC, and plasmacytoid DCs), mast cells, stress cells (marked with the expression of genes coding heat shock protein family), and *ISG15^+^*cells (**Figure 4G, H, Figure S5A**). Since tumor-associated macrophages (TAMs) play a critical role in shaping the ESCC TME^43–45^, we explored how distinct macrophage subtype may contribute to an immunosuppressive TME in relapsed tumors and observed that *SPP1^+^TREM2^+^* and *SPP1^+^* macrophages exhibited the highest immunosuppressive scores^46^ among all macrophage subtypes (**Figure 4I**). The expression of SPP1 and TREM2 in TAMs are known pro-tumorigenic factors in various cancer types^43,47–50^. Functionally, these two macrophage subtypes shared the pathways related to cell chemotaxis and migration (**Figure S4C**). We observed a significant increase in the proportion of *SPP1^+^*macrophages and *SPP1^+^TREM2^+^* macrophages in relapsed tumors compared to normal tissues and treatment-naive primary tumors (**Figure 4J, K**), contrasting with the patterns of other myeloid subtypes (**Figure S5B**). To validate these findings, we conducted an mIHC analysis on the validation cohort. This analysis confirmed a markedly higher abundance of both *SPP1^+^TREM2^+^*macrophages and *SPP1^+^* macrophages in Rel samples compared to matched Pre-Rel and Pre-NR samples (**Figure 4L, M**). These results reinforce the critical role of these macrophage subtypes in contributing to the immunosuppressive microenvironment associated with tumor relapse. Together, our results highlight the cooperative roles of NRF2-activated tumor cells and specific macrophage subtypes in driving chemoradiotherapy resistance through TME remodeling.

### NRF2-Activated Tumor Cells Exclusively Interact with SPP1^+^TREM2^+^ Macrophages in Relapsed Samples

To investigate the interactions between NRF2-activated tumor cells and the above macrophages, we analyzed cell-cell interactions within the TME using ST data integrated with scRNA-seq data. By deconvoluting the cellular composition of spatial transcriptomics spots (see **Methods**), we compared the spatial co-localization of NRF2-activated epithelial cells with *SPP1^+^* macrophages and *SPP1^+^TREM2^+^*macrophages. Our analysis revealed distinct spatial infiltration patterns for epithelial subtypes and macrophage subtypes, including *SPP1^+^* macrophages and *SPP1^+^TREM2^+^*macrophages (**Figure S6**). Specifically, NRF2-activated epithelial cells showed preferential co-localization with *SPP1^+^TREM2^+^* macrophages exclusively in relapsed samples (**Figure 5A, B**). In contrast, co-localization between NRF2-activated epithelial cells and SPP1^+^ macrophages was observed in both Pre-Rel and Rel tissues (**Figure 5C, D**). These spatial interaction patterns were further validated using mIHC analysis. Physical proximity between *SPP1^+^TREM2^+^* macrophages and AKR1B10^+^PanCK^+^ tumor cells was predominantly observed in Rel samples, with minimal interactions detected in paired Pre-Rel and Pre-NR samples (**Figure 5E, F**). These findings highlight a unique interaction between NRF2-activated tumor cells and *SPP1^+^TREM2^+^*macrophages in relapsed tumors, suggesting that these interactions play a critical role in reshaping the TME to support tumor progression and resistance to therapy.

**Figure 5.**
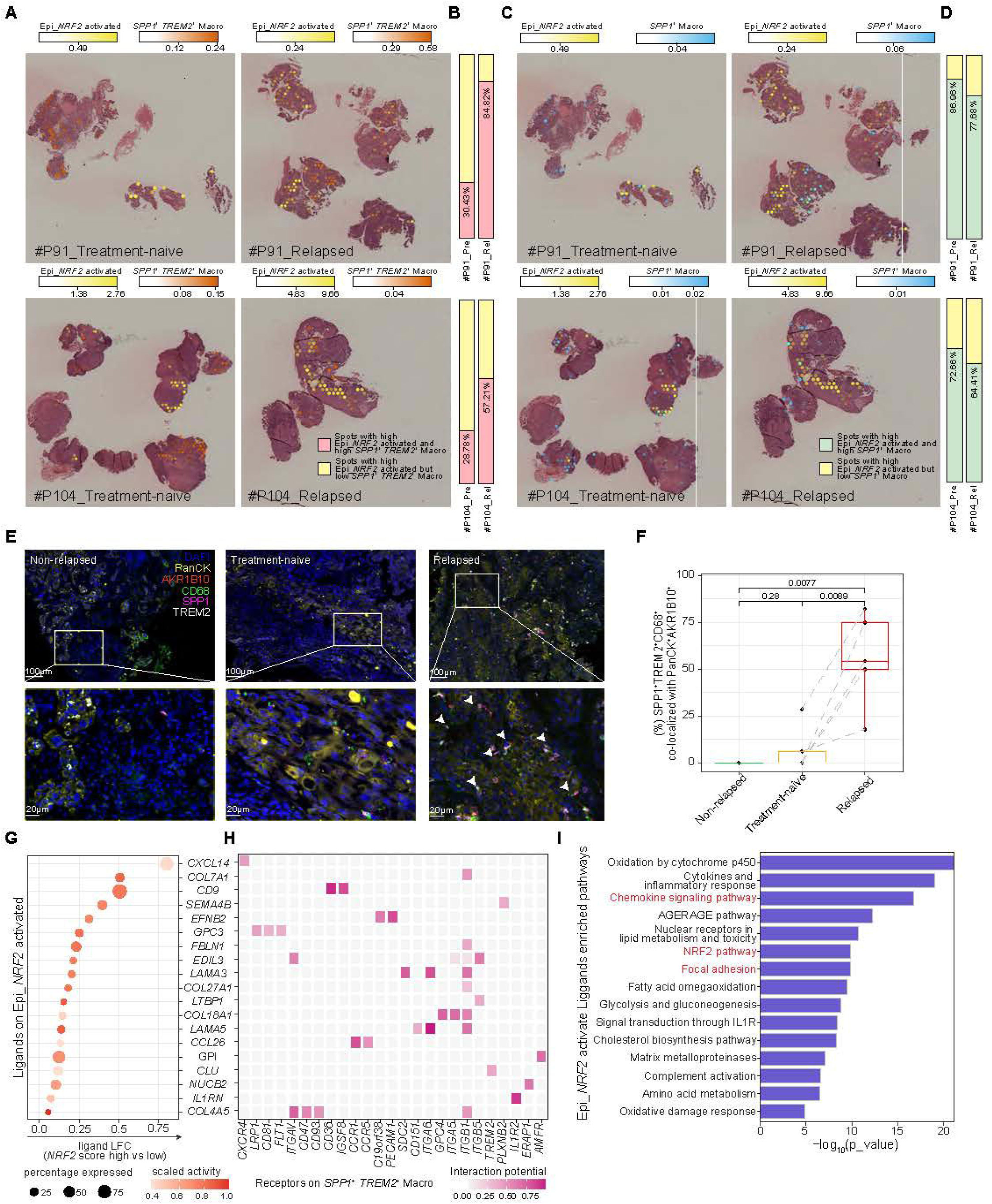
Crosstalk between NRF2-activated epithelial cells and SPP1⁺TREM2⁺ macrophages in the relapsed tumor microenvironment. **A, C**) Spatial scatter pie plots illustrating deconvolution scores of NRF2-activated epithelial cells colocalized with SPP1⁺TREM2⁺ macrophages (A) and SPP1⁺ macrophages (C) in pre-treatment (left) and paired relapsed tissues (right). The deconvolution scores represent the relative degree of RNF2 activation. **B, D**) Bar plots comparing the proportions of NRF2-activated epithelial cell score-high spots colocalized with or without SPP1⁺TREM2⁺ macrophages (B) and SPP1⁺ macrophages (D) between pre-treatment and post-treatment relapsed tissues. **E)** Representative multiplex immunohistochemistry (mIHC) images of pre-treatment non-relapsed (NR) tissues, and paired pre- and post-treatment relapsed tissues showing colocalization of SPP1⁺TREM2⁺CD68⁺ macrophages and AKR1B10⁺PanCK⁺ epithelial cells. Scale bars: upper panel, 100 µm; zoomed-in panel, 20 µm. **F)** Boxplot quantifying the percentage of SPP1⁺TREM2⁺CD68⁺ macrophages colocalized with AKR1B10⁺PanCK⁺ epithelial cells across pre-treatment NR (green, n = 5), paired pre-treatment (orange, n = 5), and post-treatment (red, n = 5) relapsed tissues in the validation cohort. Median ±1 quartile, with whiskers extending to values within 1.5× the interquartile range (IQR). P value is based on two-sided Wilcoxon signed-rank test. **G)** Top-ranked ligands predicted to regulate SPP1⁺TREM2⁺ macrophages by NRF2 pathway score-high epithelial cells, identified using NicheNet. **H)** Ligand-receptor interaction pairs highlighting crosstalk between NRF2-activated epithelial cells and SPP1⁺TREM2⁺ macrophages, ranked by ligand activity. **I)** KEGG pathway enrichment analysis showing representative pathways of potential ligand genes expressed in NRF2-activated epithelial cells. P value is based on Fisher exact ’s test. **A-D**) The two Pre-Rel and Rel sample pairs the mechanism investigation cohort.

To further explore the molecular mechanisms underlying this interaction, NicheNet analysis^51^ revealed several highly activated ligands preferentially expressed in NRF2-activated tumor cells. These included chemokines (*CXCL14* and *CCL26*) and laminins (*LAMA3* and *LAMA5*) (**Figure 5G**). Correspondingly, their receptors were predominantly expressed on *SPP1^+^TREM2^+^* macrophages, demonstrating a strong interaction potential. Key receptors included chemokine receptors (*CXCR4*, *CCR1*, and *CCR5*) and integrins (*ITGAV, ITGA6, ITGA5, ITGB1,* and *ITGB5*) (**Figure 5H**). Moreover, the downstream targets of these ligands were significantly enriched in pathways related to chemokine signaling, NRF2 activation, and focal adhesion (**Figure 5I**). These findings collectively suggest that NRF2-activated tumor cells actively recruit *SPP1*^+^*TREM2*^+^ macrophages via chemokine and adhesion-mediated pathways, thereby fostering an immunosuppressive TME. This interaction highlights a critical mechanism by which NRF2 signaling contributes to TME remodeling and tumor progression.

We further investigated the alterations of T cell states within the single-cell atlas, given their critical anti-tumor role^52,53^ role in ESCC^54–56^. This analysis encompassed 9 CD4^+^ T cell subtypes and 11 CD8^+^ T cell subtypes (**Figure S7A-F)**. Notably, naive and memory T cells exhibited reduced proportions in relapsed tumors, indicating impaired effector function in T cells among relapsed patients^57–60^. Intriguingly, the proportion of precursor exhausted CD8^+^ T cells (CD8^+^ pre-T_EX_) was significantly elevated in relapsed tissues (**Figure S7F**). CD8^+^ pre-T_EX_ cells have the potential to differentiate into exhausted T cells (T_EX_)^61^, which are marked by diminished effector function and self-renewal capacity^62^. Since T cell exhaustion is often linked to macrophage activity in the TME^63–65^, we used CellChat^66^ to infer reciprocal cell-cell interaction networks between macrophages and T cells. This analysis revealed that outgoing signaling in the TME was predominantly mediated by *SPP1^+^TREM2^+^* macrophages and *SPP1^+^* macrophages, while incoming signals were largely received by CD8^+^ T_EX_ and CD8^+^ pre-T_EX_ cells (**Figure S7G**). When considering *SPP1^+^TREM2^+^* macrophages, which demonstrated the strongest outgoing interactions with CD8^+^ T cells, we found that their interaction weights with CD8^+^ T_EX_ and CD8^+^ pre-T_EX_ were particularly high (**Figure S7H**). Additionally, in the ADGRG signaling pathway, *SPP1^+^TREM2^+^*macrophages were identified as the most important signal senders, while CD8^+^ T_EX_ and CD8^+^ pre-T_EX_ cells were the dominant signal receivers (**Figure S7I**). ADGRG, an emerging immune checkpoint, is expressed on tumor-infiltrating lymphocytes and has been shown to correlate with T cell exhaustion^67,68^. These findings provide insight into the global cell-cell interaction network between macrophages and CD8^+^ T cells within the relapsed TME. They further emphasize the role of *SPP1^+^TREM2^+^*macrophages, recruited by NRF2-activated tumor cells, as key immunosuppressive mediators: these macrophages appear to act as a “bridge,” facilitating T cell exhaustion and contributing to immune suppression during tumor relapse.

## Discussion

Chemoradiotherapy is the most widely used therapeutic approach for ESCC; however, only a small fraction of patients achieve a durable response^2,4^, underscoring an urgent and unmet clinical need. We proposes a mechanistic model in which tumor-intrinsic activation of the NRF2 pathway underlies resistance to chemoradiotherapy in ESCC (**Figure 6**). In tumor cells, frequent gain-of-function mutations in *NFE2L2* or loss-of-function mutations in its negative regulator, *KEAP1*, result in NRF2 release, nuclear translocation, and binding to ARE elements. This activates the transcription of downstream target genes involved in antioxidative responses^69^, protecting tumor cells from treatment-induced DNA damages. Moreover, NRF2-activated tumor cells recruit immunosuppressive *SPP1^+^TREM2^+^* macrophages, which further mediate the exhaustion of CD8^+^ T cells, fostering an immunosuppressive TME that supports tumor progression. These findings provide a critical foundation for developing strategies to overcome chemoradiotherapy resistance.

**Figure 6.**
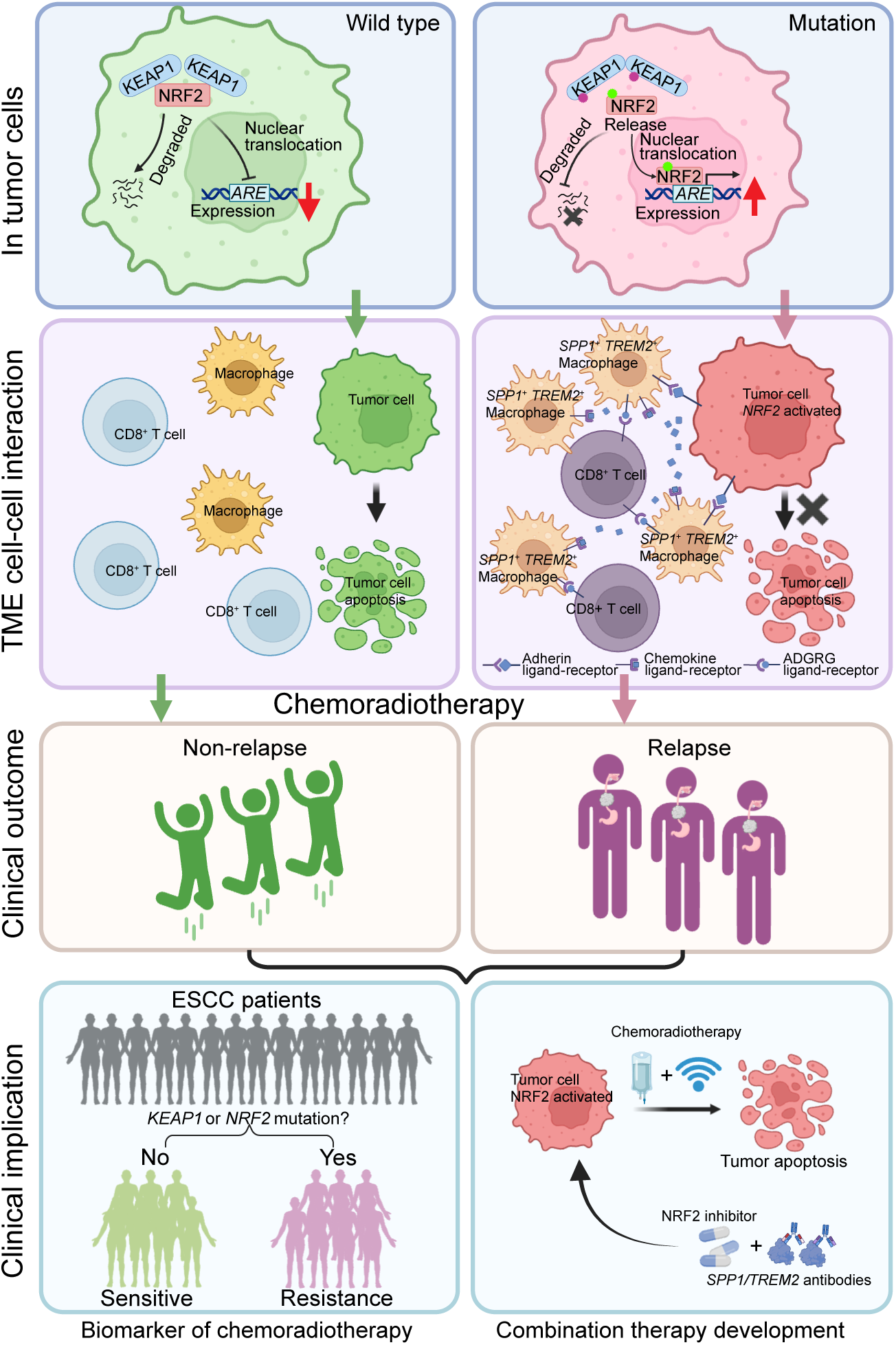
The proposed model of *NF2L2/KEAP1* mutations driving chemoradiotherapy resistance. The proposed model highlights a three-layer framework that connects molecular mechanisms to clinical implications. The left and right panels contrast tumor cells with *KEAP1/NFE2L2* wild-type versus mutation. On the right side, within tumor cells, mutated *NFE2L2* or *KEAP1* drives activation of the NRF2 pathway by stabilizing NRF2, leading to enhanced antioxidant response. In the TME, NRF2-activated tumor cells recruit SPP1⁺TREM2⁺ macrophages, which establish strong cell-cell adhesion with tumor cells. These macrophages subsequently interact with (pre-)exhausted T cells, fostering an immunosuppressive TME and promoting chemoradiotherapy resistance. In contrast, on the left side, tumor cells with wild-type *NFE2L2/KEAP1* have moderate cell-cell adhesion with *SPP1*^+^*TREM2*^+^ macrophages, thereby rendering tumor cell apoptosis. The bottom panels show *NFE2L2/KEAP1* mutations could serve as biomarkers to predict chemoradiotherapy resistance. It also underscores the potential for combination therapies targeting NRF2 pathway activation and the immunosuppressive TME to improve treatment outcomes.

Our results align with existing literature while providing novel mechanistic insights. *KEAP1* and *NFE2L2* are well-established driver genes in ESCC^70,71^ and define a distinct molecular subtype of the disease^62,64,72^. Through a comparative analysis of pretreatment samples from patients rigorously assessed in a clinical trial with balanced clinical variables, we directly link somatic alterations in these two genes to tumor relapse following chemoradiotherapy. While NRF2 activation has been reported to promote tumor-intrinsic resistance to chemo- or radiotherapy in various cancer types^24,73–75^ by multiple mechanisms, our study extends these findings by uncovering a novel role of NRF2 activation in shaping the TME. Specifically, NRF2-activated tumor cells recruit immunosuppressive *SPP1^+^TREM2^+^* macrophages that may be associated with pre-exhausted/exhausted CD8^+^ T cells, thereby creating a permissive niche for tumor relapse. While the role of *SPP1*^+^ macrophages in immunosuppression is established^65^, this key mechanistic insight into the physical interactions of *SPP1^+^TREM2^+^* macrophages with NRF2-activated tumor cells represents a significant conceptual advance. Future investigations should explore whether this mechanism exists in other tumor types.

Our findings have immediate clinical relevance. Currently, there are no approved predictive biomarkers for chemoradiotherapy response in ESCC patients. As first-line treatment options for ESCC, definitive chemoradiotherapy and surgery offer comparable survival rates^76^, leading to considerable uncertainty in clinical decision-making. Several studies^77–81^ have implicated *NFE2L2/KEAP1* in chemoradiotherapy resistance, based on evidence from cell lines, mouse models, and retrospective patient sample profiling of individual genes. While these studies have provided valuable insights, their findings do not match the depth, rigor, and comprehensive approach of this study. Our work is based on a prospective clinical trial in which patients received consistent treatment, confounding factors were well controlled, clinical responses were clearly annotated, and unbiased multi-omics characterization was performed, systematically integrating various types of genomic, transcriptomic, and TME aberration. We show that NRF2 activation drives resistance to chemoradiotherapy. However, stratifying patients based on a gene expression signature remains challenging and has seen limited success in clinical implementation. In contrast, DNA sequencing of patient tumors is well established in precision oncology. We demonstrate that nearly 40% of ESCC patients with swift relapse exhibit *NFE2L2* or *KEAP1* somatic alterations in their primary tumors, while such mutations are almost absent in non-relapsed patients. These findings provide compelling evidence that strongly supports a straightforward and effective strategy for patient stratification: direct sequencing of these two genes in pre-treatment biopsies could identify patients more likely to benefit from chemoradiotherapy. Our study also suggests new therapeutic strategies. NRF2 has emerged as a promising therapeutic target, with several inhibitors currently under clinical investigation^82^. Targeting immunosuppressive macrophages with specific agents has gained increasing attention as a promising approach in cancer therapy. Our results suggest that combining NRF2 inhibitors and/or antibodies targeting *SPP1^+^TREM2^+^*macrophages with chemoradiotherapy could improve therapeutic efficacy in ESCC patients. These findings call for immediate clinical investigation to validate the utility of the proposed predictive biomarkers and combination therapies, potentially transforming the treatment landscape for ESCC.

Our study has several limitations. First, although we analyzed a large patient cohort overall, the number of samples profiled by each individual technology remains relatively limited. Nevertheless, our well-designed, integrative approach—spanning multiple data modalities (WES, WGS, RNA-seq, scRNA-seq, spatial transcriptomics, and mIHC) and independent cohorts (discovery, mechanistic, and validation)—helps mitigate this concern. The consistent signals observed across platforms and cohorts underscore the robustness of our findings. Additional sequencing in independent patient cohorts will be valuable to further validate the generalizability of these results. Second, we did not perform direct functional experiments to establish the causal effects of *NFE2L2* mutations on oxidative stress response or the recruitment of SPP1⁺TREM2⁺ immunosuppressive macrophages. While the role of *NFE2L2* mutations in activating NRF2 signaling and promoting antioxidant responses has been well characterized in cell lines and animal models, further studies—such as co-culture systems or genetic perturbation experiments—are warranted to confirm their specific impact on macrophage recruitment and phenotype. Finally, a prospective clinical trial stratifying patients based on *NFE2L2*/KEAP1 mutation status is needed to validate their predictive value for treatment response and patient outcomes in real-world settings.

## Methods

### Patient recruitment and sample collection

This study was conducted following approval by the hospital ethics committee (ESO-Shanghai 6), with informed consent obtained from all participants. It included patients with locoregional ESCC treated with definitive concurrent chemoradiotherapy or radiotherapy alone from 2010 to 2023 at Fudan University Cancer Center (Shanghai, China), Jiangsu Province Cancer Hospital (Nanjing, China), and Yancheng Third People’s Hospital (Yancheng, China). Several elderly patients received radiotherapy alone, as they were unable to tolerate chemotherapy. Patients were divided into two groups: relapsed (Rel) and non-relapsed (NR). The relapsed group included patients with newly diagnosed locoregional ESCC treated with definitive (chemo)radiotherapy, a confirmed pathological diagnosis of squamous carcinoma prior to treatment, primary tumor recurrence within the radiotherapy gross tumor volume (GTV) within 2 years post-treatment, and a confirmed pathological diagnosis of recurrent squamous carcinoma. Tumor samples were collected via endoscopic biopsy before treatment, paired with normal peripheral blood samples obtained prior to (chemo)radiotherapy, and relapsed tumor samples collected via endoscopic biopsy or esophagectomy. The non-relapsed group included patients with newly diagnosed locoregional ESCC treated with definitive (chemo)radiotherapy, a confirmed pathological diagnosis of squamous carcinoma, and no primary tumor recurrence within the radiotherapy GTV for at least 3 years post-treatment. Pretreatment tumor samples and paired normal peripheral blood samples were collected via endoscopic biopsy before treatment.

### Sample processing

The fresh-frozen tumor tissues were snap-frozen in liquid nitrogen immediately after endoscopic biopsy or surgical resection and subsequently stored at −80°C. Formalin-fixed, paraffin-embedded (FFPE) tumor tissues were obtained from archived residual clinical specimens. Blood samples were collected prior to treatment and stored at −80°C. Fresh tumor tissues designated for single-cell RNA sequencing (scRNA-seq) were collected via endoscopic biopsy, immediately preserved in Tissue Storage Solution (Miltenyi Biotec), and processed within one hour to ensure sample integrity.

### Whole-exome sequencing

Genomic DNA was extracted from tissue specimens using the QIAamp DNA kit (Qiagen), followed by library preparation based on Illumina-recommended protocols. Briefly, 1 μg of DNA was sheared into short fragments (200–300 bp) using a Covaris S220. The fragments underwent end repair, and an adenylate blocker was added to the 3’ ends. Barcode-ligated adaptors were attached to both fragment ends, and size selection for the targeted fragments was performed using E-Gel. The selected fragments were amplified through 10 cycles of polymerase chain reaction (PCR), and the resulting mixture was purified. Whole-exome capture was conducted using the TruSeq Exome Enrichment kit (Illumina), following the manufacturer’s protocol with minor modifications. The libraries were further amplified with an additional 10 PCR cycles after hybridization with capture probes, which were incubated for 24 hours at 65°C. The validated DNA libraries were then sequenced on the Illumina HiSeq 2500 sequencing system.

### WES data analysis

Read pairs in FASTQ format were trimmed and filtered for quality using fastq-mcf (https://github.com/ExpressionAnalysis/ea-utils). High-quality reads were aligned to the human reference genome (GRCh37) using the Burrows-Wheeler Aligner (BWA 0.7.12). The resulting BAM files were processed using the Genome Analysis Toolkit (GATK) to enhance alignment accuracy. Key steps included marking duplicates, performing local realignment around high-confidence insertions and deletions (INDELs), and recalibrating base quality scores. Somatic point mutations were identified using five widely-used variant callers: Muse^83^, MuTect2^84^, SomaticSniper, Radia^85^, and VarScan2^86^; NDELs were detected using PINDEL^87^, VarScan2 and Mutect2. Mutations (both point mutations and INDELs) reported by at least two callers were retained for further analyses. A multi-step filtering process was applied to ensure high-quality mutation calls, including criteria such as minimum read depth, strand-bias checks, and exclusion of artifacts using a panel of normals^88^. This pipeline ensured a robust and reliable set of somatic mutation data for downstream analyses.

Copy Number Variation and Clonality Analysis: Sequenza was used to estimate SCNAs, ploidy, and tumor purity for each tumor sample, with default settings applied. SCNA profiles outputted by Sequenza, further adjusted by sample ploidy, were analyzed using GISTIC to identify significant amplification and deletion peaks. Genome doubling status was inferred using EstimateClonality (v1.0) with input derived from Sequenza’s SCNA profiles. Mutation Signature and Timing Analysis: The R package mutationalPatterns was used to estimate the contribution proportions of mutational signatures for each tumor sample. Relative mutation timing was inferred using the MutationTimeR^89^ package, utilizing variant allele frequency, tumor purity, and SCNA profiles as inputs. Clustering and Clonal Evolution Analysis: Variant allele frequency, tumor purity, and SCNA profiles were processed with PyClone to estimate mutation clusters and calculate adjusted cancer cell fractions (CCF). The MATH score, quantifying intratumor heterogeneity, was calculated as MATH=100×MAD/median CCF, where MAD is the median absolute deviation of CCF. Cancer cell fractions from PyClone were further analyzed with sciClone to generate refined mutation clusters. Clonal evolution trajectories were inferred using clonEvol and visualized with Fishplot. The 3D bubble plots, TEDGs, and patient clustering on the moduli space were generated using CELLO^90^. CCFs given by PyClone were further provided to sciClone to generate new mutation clusters. Clonal evolution trajectories were inferred using clonEvol and visualized using Fishplot. Somatic mutation data was uploaded to mutationMapper to visualize their distribution on corresponding protein. TEDG was constructed using the mutDirectedGraph function.

### Whole-genome sequencing

DNA libraries were prepared using the TruSeq Nano DNA LT Prep Kit (Illumina), and whole-genome sequencing was performed on the Illumina NovaSeq 6000 platform. Fragment overhangs were repaired using End Repair Mix, followed by purification with AMPure XP Beads (Beckman). The purified DNA fragments underwent adenylation with A Tailing Mix and were subsequently ligated with adapters using DNA ligase. Adapter-ligated fragments were amplified and purified, and the resulting libraries were quantified using the Qubit dsDNA HS Assay (Thermo Fisher Scientific). The size distribution of sequencing libraries was assessed using the Agilent BioAnalyzer 2100 (Agilent). Paired-end reads were generated, targeting a sequencing depth of 3–5× per sample, enabling low-coverage genome-wide profiling.

### WGS data analysis

Similar to the WES alignment, read pairs in FASTQ format were trimmed and filtered with fastq-mcf (https://github.com/ExpressionAnalysis/ea-utils). The resulting high-quality reads were aligned to the human reference genome (GRCh37) using Burrows-Wheeler Aligner (BWA 0.7.12). The aligned BAM files were processed with the Genome Analysis Toolkit to enhance alignment accuracy. Key processing steps included marking duplicates, local realignment around high-confidence insertion and deletions, and base quality recalibration. SCNAs were detected using CNVkit^91,92^ following the authors’ guidelines. The analysis was conducted with default parameters and restricted to WGS mappable regions and the captured regions.

### Bulk RNA sequencing

Total RNA was isolated using the RNeasy Mini Kit (Qiagen, Germany). Strand-specific libraries were prepared using the TruSeq Stranded Total RNA Sample Preparation Kit (Illumina, USA) following the manufacturer’s protocol. Briefly, mRNA was enriched using oligo(dT) beads and fragmented into small pieces at 86°C for 6 minutes with divalent cations. First-strand cDNA synthesis was performed using reverse transcriptase and random primers, followed by second-strand cDNA synthesis with DNA Polymerase I and RNase H. The resulting cDNA fragments underwent end repair, the addition of a single ’A’ base, and adapter ligation. The final cDNA libraries were purified and enriched by PCR, quantified using a Qubit 2.0 Fluorometer (Life Technologies, USA), and validated using the Agilent 2100 BioAnalyzer (Agilent Technologies, USA) to confirm insert size and determine molar concentration. Libraries were diluted to 10 pM for cluster generation using cBot and sequenced on the Illumina NovaSeq 6000 (Illumina, USA). Library construction and sequencing were performed at the Shanghai Biotechnology Corporation, ensuring high-quality data generation.

### RNA-seq data analysis

For each sample, 33–95 million RNA-seq clean reads were obtained and aligned to the GRCh38 reference genome (ftp://ftp.ensemblgenomes.org/) using HISAT2^93^ (v2.0.477). Sequencing read counts were calculated using Stringtie^94^ (v.1.3.0) and expression levels were normalized across samples using the Trimmed Mean of M values (TMM) method. Normalized expression levels were converted to FPKM (Fragments Per Kilobase of transcript per Million mapped fragments). Differential gene expression analysis was performed using the R package *edgeR*^95^. P-values were calculated, and multiple hypothesis testing was controlled by the Benjamini-Hochberg algorithm to calculate the False Discovery Rate (FDR), with corrected P-values referred to as q-values. Differentially expressed genes (DEGs) were defined as transcripts with a fold change in expression (based on FPKM values) greater than 2.0 and a q-value less than 0.05. Gene Ontology (GO) enrichment analysis was performed using the R package *clusterProfiler*^96^, with enrichment criteria set to q-value < 0.05. Heatmaps of specific genes were generated using the R package *pheatmap*.

### Cell line radiation-sensitivity analysis

Radiation-sensitivity data, measured as AUC scores, for 533 cell lines was obtained from a published study^29^. Somatic mutation data for these cell lines was sourced from CCLE^97^. For each cell line, the NRF2 pathway score was calculated based on the expression levels of NRF2 pathway-related signatures. Cell lines were then grouped into two categories—NRF2 pathway high and NRF2 pathway low—based on their pathway scores. Survival scores (AUC values) between the two groups were compared using the Wilcoxon test, enabling an evaluation of the association between NRF2 pathway activation and radiation sensitivity across diverse cell lines.

### scRNA-seq library preparation and sequencing

Tissues were transported in a sterile culture dish containing 10 mL of 1× Dulbecco’s Phosphate-Buffered Saline (DPBS; Thermo Fisher, Cat. no. 14190144) on ice to remove residual tissue storage solution. The tissues were minced on ice and digested using 0.25% Trypsin (Thermo Fisher, Cat. no. 25200-072) and 10 µg/mL DNase I (Sigma, Cat. no. 11284932001) dissolved in PBS with 5% Fetal Bovine Serum (FBS; Thermo Fisher, Cat. no. SV30087.02). The digestion process was conducted at 37°C with shaking at 50 r.p.m. for approximately 40 minutes, with dissociated cells collected every 20 minutes to maximize cell yield and viability. The resulting cell suspensions were filtered through a 40 μm nylon cell strainer, and red blood cells were removed using 1× Red Blood Cell Lysis Solution (Thermo Fisher, Cat. no. 00-4333-57). Dissociated cells were washed with 1× DPBS containing 2% FBS. Cell viability was assessed with 0.4% Trypan Blue (Thermo Fisher, Cat. no. 14190144) using the Countess® II Automated Cell Counter (Thermo Fisher). Freshly dissociated single-cell suspensions were processed immediately according to the manufacturer’s protocol for the 10× Single Cell 3’ v3 Kit (10× Genomics, Pleasanton, CA). Library preparation and sequencing were performed on the NovaSeq 6000 platform, ensuring high-quality single-cell RNA sequencing data.

### scRNA-seq data analysis

Raw reads from scRNA-seq were processed using the Cell Ranger pipeline (version 5.0.0, 10× Genomics) and mapped to the GRCh38 reference genome to generate gene count matrices indexed by cell barcodes. The resulting gene-barcode matrices were analyzed using the Seurat R package (version 4.3.2). Cells expressing at least 200 genes and genes detected in at least 3 cells were retained for further analysis (**Table S2**). All samples were merged into a single Seurat object and filtered based on the following quality control criteria: nFeature_RNA between 400 and 7500, nCount_RNA between 800 and 10,000, and mitochondrial gene content below 30%. These preprocessing steps ensured the retention of high-quality cells for downstream analyses.

Clustering and dimension reduction were performed on the filtered Seurat object, which was normalized and scaled with the mitochondrial gene percentage regressed out. The top 2000 most variable genes were identified using the “FindVariableFeatures” function in Seurat (v4.3.2) and used for principal component analysis (PCA). Nearest neighbors were identified using the “FindNeighbors” function, and graph-based clustering was performed with “FindCluster” to define cell subtypes. The uniform manifold approximation and projection (UMAP) algorithm was applied for cell subtype visualization. To correct for batch effects, the harmony algorithm (R package *Harmony*^98^) was applied before clustering. Cells were first partitioned into broad categories: epithelial cells, myeloid cells, T and NK cells, B and plasma cells, endothelial cells, and fibroblasts. These clusters were further subclustered into major subtypes for each category. Cluster identities were assigned based on marker gene expression and manually reviewed to ensure accurate cell type annotation. Differentially expressed genes (DEGs) among clusters were identified using the “FindAllMarkers” function in Seurat. The non-parametric Wilcoxon rank-sum test was applied to calculate p-values for comparisons across clusters. To control for multiple testing, adjusted p-values for all genes in the dataset were computed using the Bonferroni correction, ensuring robust identification of significant DEGs. The NRF2 pathway activation score was calculated using NRF2 pathway genes from the WIKIPATHWAYS gene sets downloaded from the Molecular Signatures Database^99^ (MsigDB, https://www.gsea-msigdb.org/gsea/msigdb/). For single-cell analysis, the gene signature in each cell was scored using the “enrichIt” function in the escape R package (v1.12.0), with the “ssGSEA” method and groups set to 1000. For the NRF2 activation signature score, a gene list from Härkönen et al.^42^ was used. The “AddModuleScore” function in the Seurat package was applied to compute the NRF2 activation signature score, ensuring accurate quantification of pathway activation at the cellular level.

### Spatial transcriptomics sequencing

Freshly prepared tissue samples were processed following the 10× Spatial Transcriptomics protocol (10× Genomics, Pleasanton, CA). After tissue fixation, hematoxylin and eosin (HE) staining and imaging were conducted. Permeabilization of selected tissues was performed in accordance with the 10× Spatial 3’ v3 kit protocol. Library preparation and sequencing were carried out on the Illumina platform, using spatially barcoded mRNA-binding oligonucleotides as specified in the default protocol by 10× Genomics.

### ST data analysis

Raw sequencing data from spatial transcriptomics were quality-checked and mapped using Spaceranger. The resulting gene-spot matrices were analyzed in Seurat (version 4.3.2) in R. Quality control of spatial transcriptomic data was performed using parameters including total spots, median UMIs/spot, median genes/spot, and median mitochondrial genes/spot (Table S3). Spots retained for subsequent analyses met the following criteria: a minimum detected gene count of 200 genes, and genes expressed in fewer than three spots were excluded (Table S3). To improve the mapping of scRNA-seq clusters to spatial spots, the cellular composition of each spatial spot was deconvoluted using the cell2location Python module (v0.1.3), following the tutorial guidelines from the Cell2location website (https://cell2location.readthedocs.io/en/latest/notebooks/cell2location_tutorial.html). The single-cell regression model was trained with parameters max_epochs=250 and lr=0.002, while the final cell2location model was trained with parameters max_epochs=30,000. This integration provided a detailed spatial map of cellular compositions.

### ST cell-To-cell communication analysis

First, we identified NRF2-activated epithelial cells, SPP1⁺TREM2⁺ macrophages, and SPP1⁺ macrophages in the spatial transcriptomics (ST) data based on their deconvolution scores. Spatial spots with a cell-type deconvolution score higher than the median value for that cell type on each ST slides were classified as cell-type infiltrated spots. A spatial spot containing both NRF2-activated epithelial cells and SPP1⁺TREM2⁺ macrophages was labeled as NRF2-activated epithelial cell and SPP1⁺TREM2⁺ macrophage co-localization. Similarly, a spatial spot containing both NRF2-activated epithelial cells and SPP1⁺ macrophages was labeled as NRF2-activated epithelial cell and SPP1⁺ macrophage co-localization.

Next, we used *NicheNet*^51^ package to infer the interaction mechanisms between NRF2-activated epithelial cells (sender cells) and SPP1⁺TREM2⁺ macrophages (affected cells) in the scRNA-seq dataset. For ligand-receptor interactions, only clustered cells with gene expression levels above 10% were considered. NRF2-activated cells with NRF2 pathway scores higher than the median were classified as the NRF2 pathway score high group, while others were classified as the low group. The NRF2 pathway score high group served as the condition of interest for prioritizing ligands. The top 300 ligands and top 2000 target genes from differentially expressed genes of the sender and affected cells were extracted for paired ligand-receptor activity analysis. To assess functional gene sets enriched by top-ranked ligands, gene sets from the Kyoto Encyclopedia of Genes and Genomes (KEGG) database, downloaded from MsigDB^99^, were used as potential targets. Functional enrichment was cross-validated using the Fisher’s exact test in two rounds, and the average p-value was calculated to ensure robust prioritization of ligand-receptor interactions and their functional implications.

Finally, we used *CellChat*^66^ (version 2.1.2) to analyze cell-to-cell communication between macrophages and T cells. A CellChat object was created by grouping the cells into predefined clusters. The ligand-receptor interaction database used for this analysis was “CellChatDB.human”, without additional supplementation. Preprocessing steps were conducted using default parameters provided by the package. To infer the communication network, the functions “computeCommunProb” and “computeCommunProbPathway” were applied for each ligand-receptor pair and each signaling pathway, separately.

### Multiplex immunohistochemistry (mIHC) staining

For mIHC staining, formalin-fixed paraffin-embedded (FFPE) tissue sections were cut to a thickness of 5 μm and melted at 60°C for 120 minutes. Sections were dewaxed in three changes of xylene (5 minutes each) and rehydrated in 100% ethanol (3 × 10 minutes) and 95% ethanol (10 minutes). Antigen retrieval was performed at 98°C for 10 minutes in 10 mM citrate buffer (pH 6). Endogenous peroxidase activity was quenched by incubating sections in 3% hydrogen peroxide for 10 minutes. After cooling for 30 minutes, sections were washed three times (5 minutes each) in 0.02% tris-buffered saline with Tween 20 (TBST) under gentle agitation. Tissue sections were incubated overnight at 4°C with the following primary antibodies: DAPI, anti-CD68, anti-SPP1, anti-TREM2, anti-PanCK, anti-AKR1B10, and anti-CD8. Following primary antibody incubation, sections were washed three times (3 minutes each) in 0.02% TBST and incubated with SignalStain® Boost IHC Detection Reagent specific to the species of the primary antibody. Signal amplification was performed using TSA Plus working solution diluted 1:50 in 1× amplification diluent, applied at room temperature for 10 minutes as per the manufacturer’s instructions. Sections were then washed five times (5 minutes each) in 0.02% TBST, mounted with ProLong Gold Anti-Fade Reagent (Life Technologies), and imaged using a Zeiss confocal microscope. For quantification of mIHC images, QuPath (version 0.5.1) was used. Cells in mIHC images were detected using DAPI staining and annotated based on fluorescence intensity. To identify co-localized cells, SPP1⁺TREM2⁺CD68⁺ cells were selected, and the Manhattan distance was calculated between these cells and all others in the mIHC image. For each SPP1⁺TREM2⁺CD68⁺ cell, the 10 closest cells were identified. If AKR1B10⁺PanCK⁺ tumor cells were among these, the SPP1⁺TREM2⁺CD68⁺ cell was recorded as co-localized with AKR1B10⁺PanCK⁺ tumor cells.

### Cell culture

ECA109 human esophageal cancer cells were cultured in DMEM medium (Gibco, USA) supplemented with 10% fetal bovine serum (FBS, Gibco, USA) and 1% Penicillin-Streptomycin solution (Pen-Strep, Gibco, USA). Cells were maintained in a humidified incubator at 37°C with 5% CO₂ to ensure optimal growth conditions.

### Western blotting

ECA109 cells were cultured in 24-well plates (Nest Biotechnology) and transfected with plasmids using a transfection reagent. After 48 hours, the cells were lysed using RIPA buffer to extract proteins. Proteins were separated on a 10% SDS-PAGE gel and transferred to PVDF membranes. The membranes were blocked with 5% non-fat milk dissolved in TBST, followed by sequential incubation with primary antibodies and HRP-conjugated secondary antibodies. Protein bands were visualized using the ECL detection kit (Sigma Aldrich, USA) and imaged with the GelView 6000Plus multi-function imaging system (BLT Photon Technology, China).

### Firefly luciferase reporter assay

ECA109 cells were cultured in 24-well plates (Nest Biotechnology) and transfected with the ARE firefly luciferase plasmid, Renilla luciferase plasmid, and the Nrf2 plasmid containing common mutations, using a transfection reagent. After 24 hours of transfection, the cells were stimulated with or without 40 nM PMA and 1 μM ionomycin for 7 hours. Firefly and Renilla luciferase activities were measured using the Dual-Glo® Luciferase Assay System (Promega, USA) according to the manufacturer’s instructions. Each experiment was performed with at least three replicates to ensure reliability and reproducibility of the results.

### Statistical Analysis

All statistical analyses were performed using R (version 4.2.1). Continuous variables were compared using the Student’s t-test or Wilcoxon rank-sum test for normally and non-normally distributed data, respectively. Categorical variables were analyzed using the Chi-square test or Fisher’s exact test. P < 0.05 is considered statistically significant

## Data and Code Availability

The newly generated raw WES/WGS data and bulk RNA-seq data have been submitted to Genome Sequence Archive (https://ngdc.cncb.ac.cn/gsa). The processed data including mutation data, RNA-seq expression data, scRNA-seq data and ST data have been deposited in Mendeley Data (989mv6y5vh). Code used for all processing and analysis is available upon request.

## Acknowledgments

This study was supported by was supported by the National Natural Science Foundation of China (U22A20326, 81872454, and 82003033), and Shanghai Rising-Star Program (23YF1406700) and Barnhart Family Distinguished Professorship in Targeted Therapies for the University of Texas MD Anderson Cancer Center.

## Author contributions

K.Z. and H.L. conceived and designed the study, J.D. led sample collection and characterization, Z.X. and H.C. led data analysis, Y. C., Y.L., W. L., W.C., J.C., J.Y., X.G., Q.F., J.S., Q.C., Y.L. F.D., M.S., Y.Z., K.C., Z.Z., Y.H., W.Z., J.L., Z.X., H.Z., Q.L., D.A., Y.C. J.L., contributed to investigation, J.D. Z.X. H.C., K.Z., and H.L. wrote the manuscript.

## Competing interests

H.L. is a shareholder and scientific advisor of Precision Scientific Ltd.

