## Supplementary figures for "NRF2 pathway activation and SPP1⁺TREM2⁺ macrophages drive chemoradiotherapy resistance in esophageal squamous cell carcinoma"

#### Supplementary Figure 1 related to Figure 1

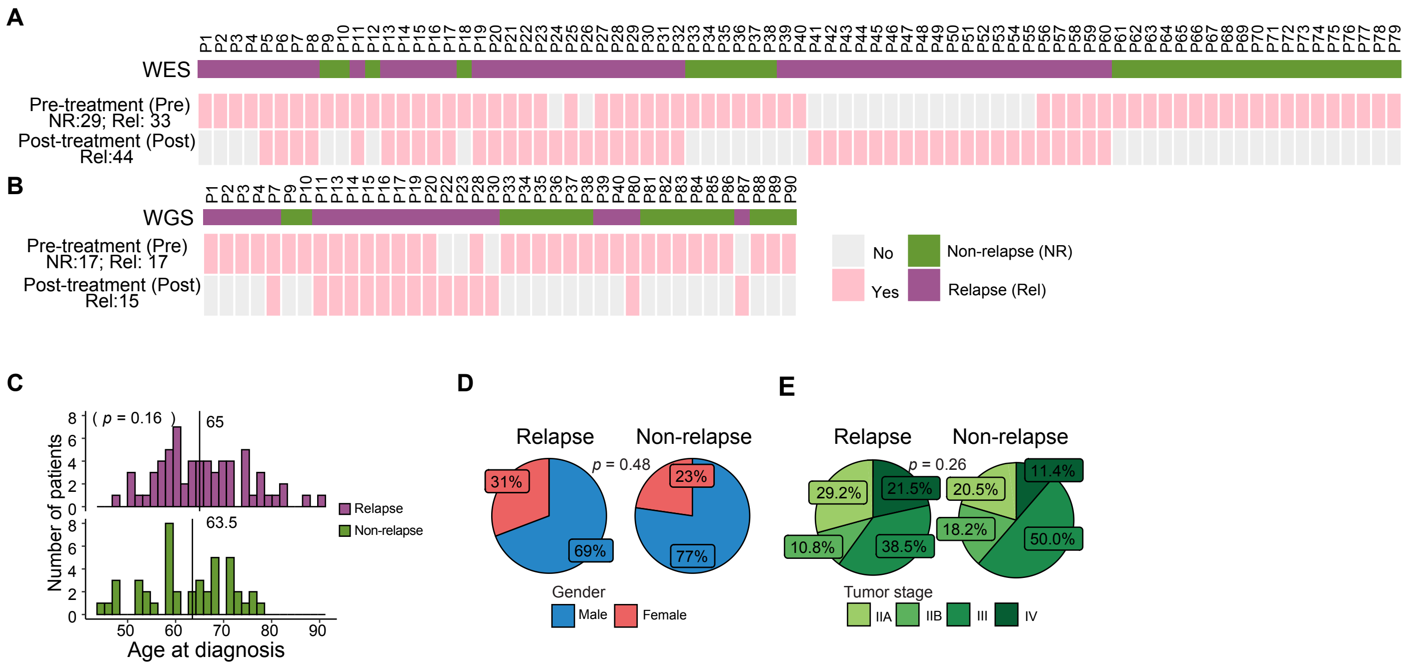


**Fig. S1. Clinical characteristics of discovery cohort.**

**(A-B)** Heatmap showing the sample compensation of the WES (A) and WGS (B) sequencing.

**(C)** Bar plots showing the age at diagnosis of relapse (upper panel) and non-relapse (bottom) patients. (D-E) Pie plots comparing the proportions of gender (D) and tumor stages (E) between relapsed and non-relapsed ESCC patients.

#### Supplementary Figure 2 related to Figure 2

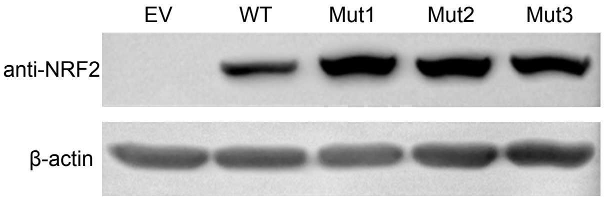


**Fig. S2. *NFE2L2* mutation influences the expression of NRF2.**
Western blot showing the expression of NRF2 of different NFE2L2 mutation sites.

#### Supplementary Figure 3 related to Figure 3

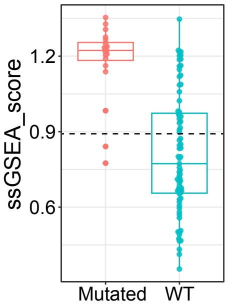


**Fig. S3. NRF2 pathway activation may not depend on *NFE2L2/KEAP1* mutations.** Box plot showing the ssGSEA score of NRF2 pathway in TCGA ESCC patients, stratified by the mutation status of NFE2L2/KEAP1.

#### Supplementary figure 4 related to Figure 4

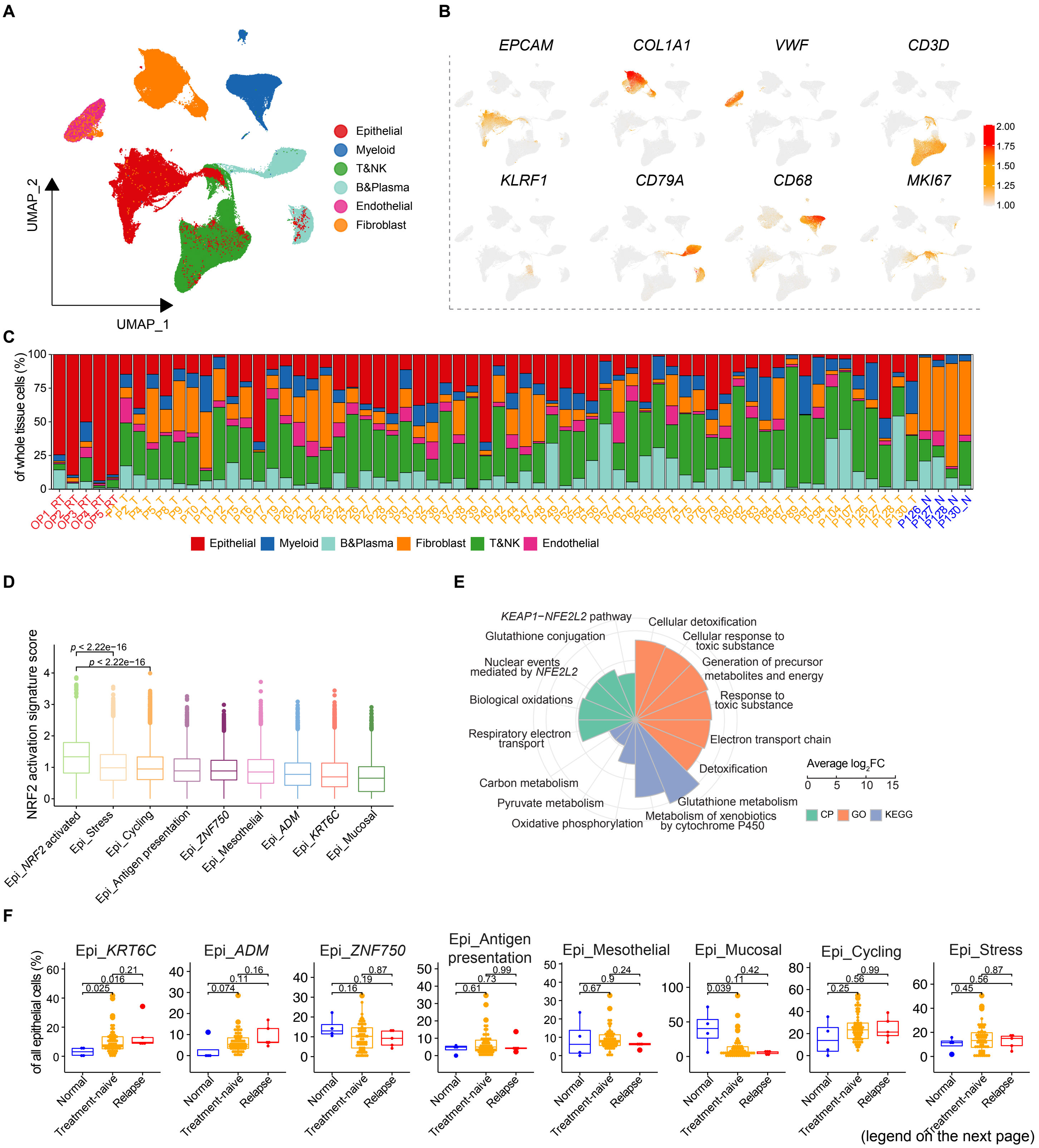


(legend on the next page)

**Fig. S4. Single-cell transcriptomic atlas of normal/pre-treatment and relapsed tissues in ESCC patients.**

**(A)** UMAP plot showing the main types of the whole tissue cells obtained from normal/pre-treatment and relapsed tissues of ESCC patients. Each dot represents one cells. Colors of cells are coded accroding to cell types.

**(B)** Feature plots of the expression of marker genes used to annotate main types.

**(C)** Bar plots showing the percentage of each main type in the whole tissue.

**(D)** Box plot showing the signature socre of NRF2 activation signature score among all epithelial sub-clusters.

**(E)** Enrichment of the differential expressed genes in NRF2 activated epithelial in CP, GO, and KEGG pathways.

**(F)** Difference in the epithelial sub-clusters' proportion of all epithelial cells,among adjacent normal, treatment-naive, and relapsed tumor tissues.

CP: canonical pathways, KEGG: Kyoto Encyclopedia of Genes and Genomes. GO: Gene Ontology. A two-sided Wilcoxon signed-rank test was used to assess the statisticval difference in (d) and (f).

### Supplementary figure 5 related to Figure 4

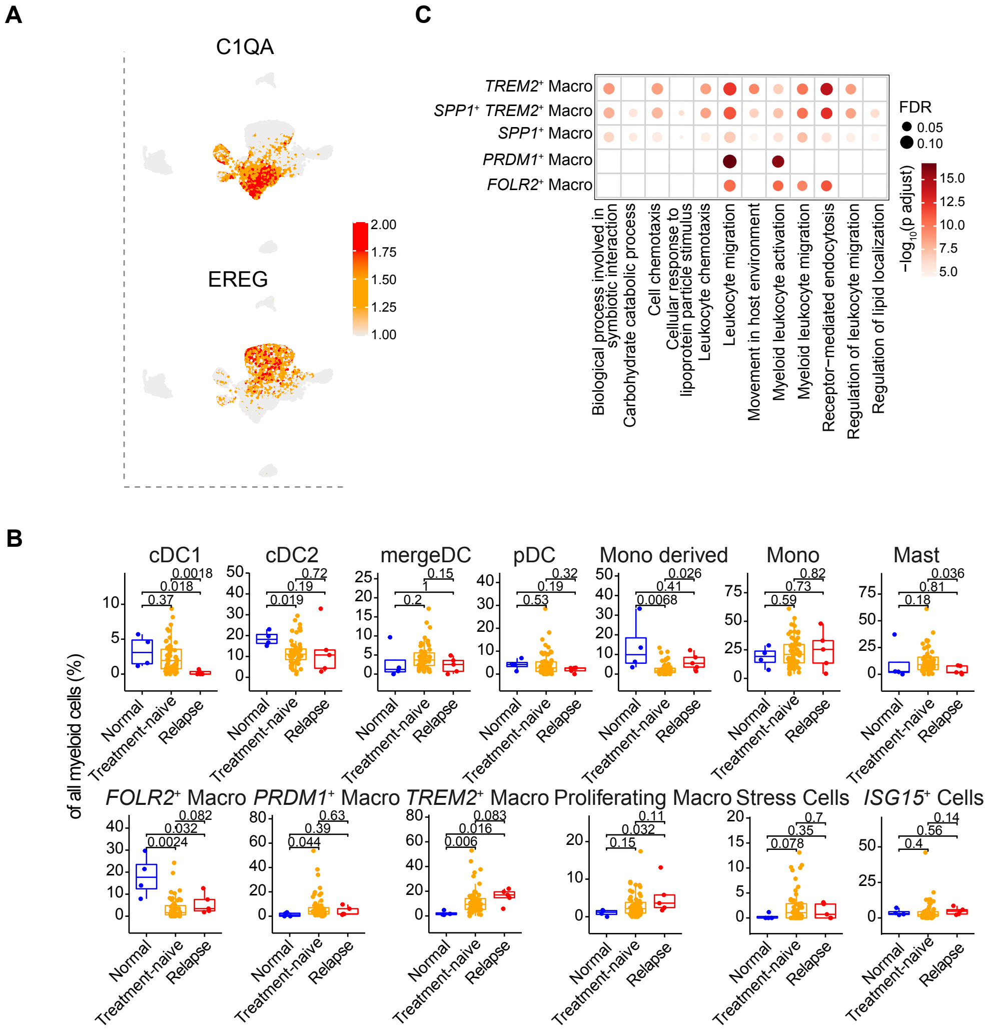


**Fig. S5 Characterization of myeloid cells, related to Figure 4**

**(A)** Feature plots of the expression of marker genes used to differ macrophages and monocytes.

**(B)** Difference in the myeloid sub-clusters' proportion of all epithelial cells,among adjacent normal, treatment-naive, and relapsed tumor tissues.

**(C)** Dot plots showed the enrichment of macrophage differentially expressed genes in GO pathways.

A two-sided Wilcoxon signed-rank test was used to assess the statisticval difference in (c).

### Supplementary figure 6 - related to figure 5

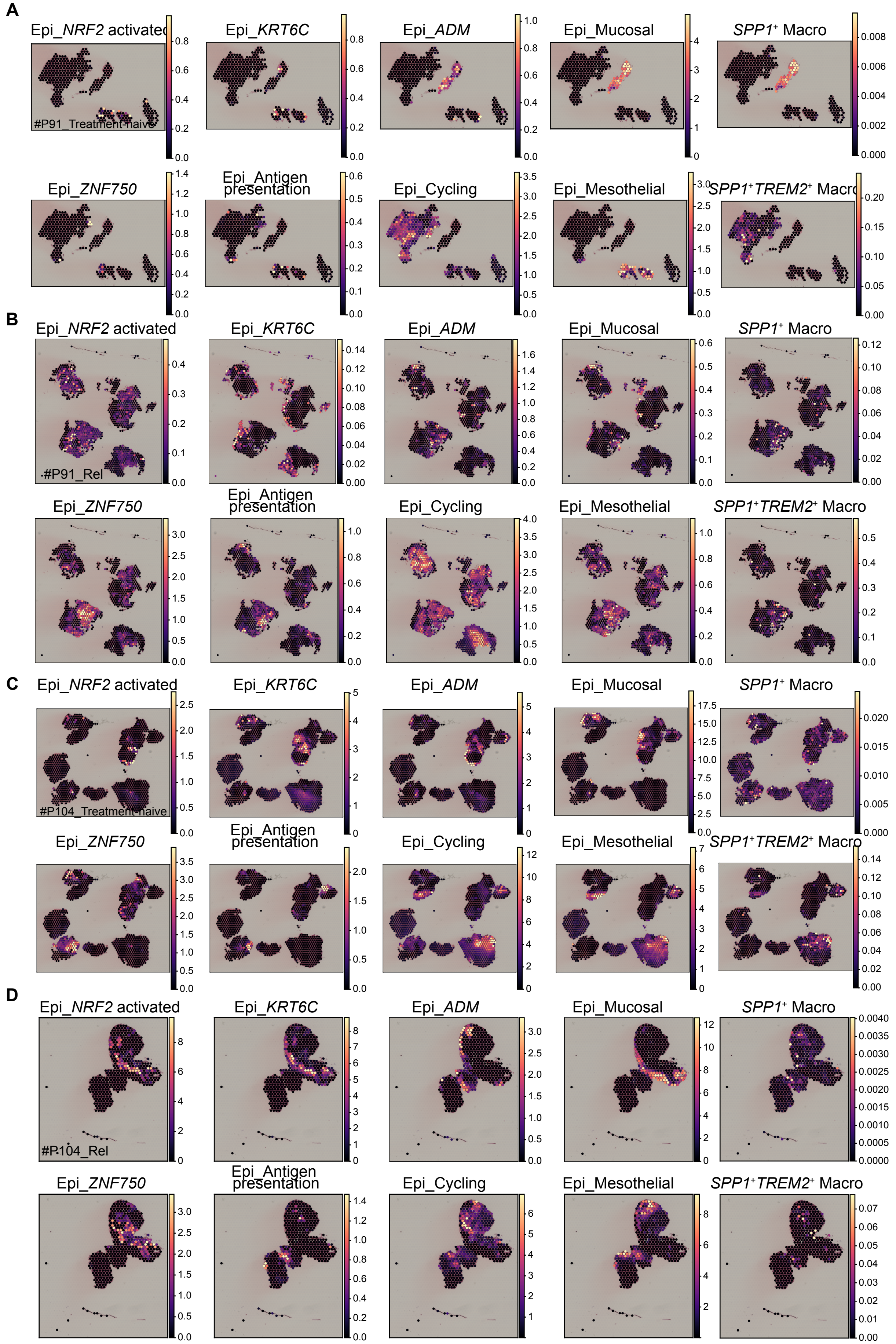


(legend on the next page)

**Fig. S6. Spatial infiltartion score of epithelial sub-clusters, *SPP1*⁺ macrophages, and *SPP1*⁺*TREM2*⁺ macrophages, related to Figure 5.**

**(A-D)** Spatial Feature plots showing the deconvolution score of epithelial sub-clusters, *SPP1*⁺ macrophages, and *SPP1*⁺*TREM2*⁺ macrophages in pre- and post-treatment relapsed tissues respectively.

#### Supplementary figure 7

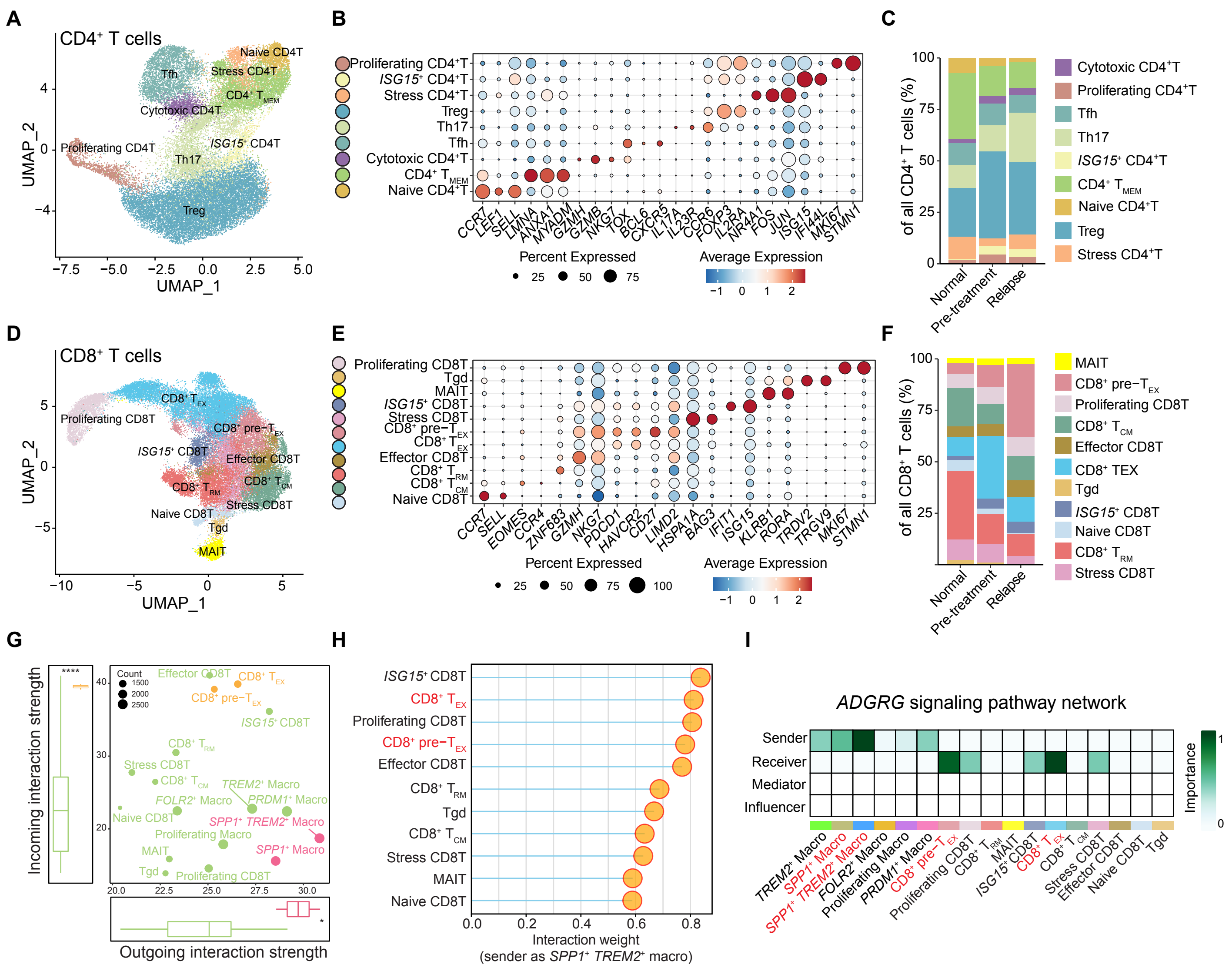

**Fig. S7. Characteristics of T cell subsets and cell-cell interaction with macrophages**

**(A,D)** UMAP plot showing the subtypes of the CD4 (**A**) and CD8 (**D**) T cells obtained from normal/pre-treatment and relapsed tissues of ESCC patients. Each dot represents one cells. Colors of cells are coded accroding to cell types.

**(B,E)** Dot plots showed the averayg expresstion (color) and expression proportion (size) of markers in subcluster of CD4 (**B**) and CD8 (**E**) T cells.

**(C,F)** Bar plots showing the differences in the percentage of CD4 (**C**) and CD8 (**F**) subtypes among normal (N), primary tumor (PT), and relapsed tumor tissues (RT)

**(G)** The potentail outgoing and incoming interaction strength of each macrophage and CD8$^{+}$ T cell cluster by CellChat analysis.

**(H)** Lolipop plot showing the interaction weight from *SPP1*$^{+}$*TREM2*$^{+}$ macrophages to CD8$^{+}$ T cell subtypes.

**(I)** Heatmap showing the predicted interaction potential between macrophage and CD8$^{+}$ T cell subtypes in the ADGRG signaling pathway.
